# Arf1-PI4KIIIβ positive vesicles regulate phosphatidylinositol-3-phosphate signalling to facilitate the fission of lysosomal tubules

**DOI:** 10.1101/2023.03.24.534147

**Authors:** Maxime Boutry, Laura F. DiGiovanni, Nicholas Demers, Aaron Fountain, Sami Mamand, Roberto J. Botelho, Peter K. Kim

**Author notes:** Corresponding author: Peter K. Kim Peter Gilgan Centre for Research & Learning The Hospital for Sick Children 686 Bay Street, 19-9800 Toronto, Ontario M5G0A4, Canada (P.K.K).

## Abstract

Formation and fission of tubules from lysosomal organelles, such as autolysosomes, endolysosomes or phagolysosomes, are required for lysosome reformation. However, the mechanisms governing these processes in the different forms of lysosomal organelles are poorly understood. For instance, the role of phosphatidylinositol-4-phosphate (PI(4)P) is unclear as it was shown to promote the formation of tubules from phagolysosomes but was proposed to inhibit tubule formation on autolysosomes because the loss of PI4KIIIβ causes extensive lysosomal tubulations. Using super-resolution live-cell imaging, we show that Arf1-PI4KIIIβ positive vesicles are recruited to tubule fission sites from autolysosomes, endolysosomes and phagolysosomes. Moreover, we show that PI(4)P is required to form autolysosomal tubules and that increased lysosomal tubulation caused by loss of PI4KIIIβ represents impaired tubule fission. At the site of fission, we propose that Arf1-PI4KIIIβ positive vesicles mediate a PI(3)P signal on lysosomes in a process requiring the lipid transfer protein SEC14L2. Our findings indicate that Arf1-PI4KIIIβ positive vesicles and their regulation of PI(3)P are critical components of the lysosomal tubule fission machinery.

## Introduction

Lysosomes are degradative organelles and metabolic signaling hubs that play an essential role in cellular homeostasis^1,2^. Dysfunction of lysosomes has been linked to many human diseases ranging from neurodegeneration to metabolic disorders^3,4^, highlighting the importance of lysosomal functions for human health. Lysosomes are acidic organelles containing catabolic enzymes enabling the digestion of macromolecules such as proteins, lipids, or carbohydrates. They receive cargos from several pathways, including endocytosis, phagocytosis and autophagy, in a fusion-dependent manner, generating endolysosomes, phagolysosomes and autolysosomes, respectively^1^. These parent lysosomal organelles are highly dynamic, undergoing various membrane fission events that occur through vesiculation, splitting or membrane tubulation^5^. In particular, the formation and fission of tubules from the parent lysosomal organelles are needed for recycling lysosomal membrane components to reform competent lysosomes^1,6–8^. This is vital for regenerating functional lysosomes that were consumed in the formation of the parent lysosomal organelles. Defects in proteins involved in tubulation and fission are associated with familial forms of neurodegenerative disease, such as Hereditary Spactic Paraplegias and Parkinson’s disease, suggesting that lysosomal tubulation is crucial to cellular homeostasis^9,10^. However, the mechanisms for lysosomal tubule formation and fission are still incompletely understood^5^.

Lysosome reformation by tubulation can be divided into three general steps^5^: the budding step that forms the nascent tubule^11^, the elongation step, and the fission of the tubule, which involves constriction and scission machinery, including dynamins^12^. Although these three steps are shared among the parent lysosomal organelles, it is not clear whether they also share the same molecular machinery or whether specific components exist for each different parent lysosomal organelles^1^.

Phosphoinositides, a group of phospholipids with various degrees of phosphorylation on the inositol headgroup of phosphatidylinositol, act at different steps on the tubulation-mediated recycling of the lysosomal membrane. For instance, phosphatidylinositol-4,5-biphosphate (PI(4,5)P_2_) was shown to mediate tubule extension and the recruitment of the scission machinery for autolysosomes^5,11^. Additionally, PI(4)P was described to inhibit the formation of tubules from autolysosomes because depletion of PI4KIIIβ (phosphatidylinositol 4-kinase IIIβ)—an enzyme producing PI(4)P—led to extensive tubulation of lysosomes^13^. This proposed function of PI(4)P differs from recent studies on both phagolysosomes^14^ and endosomes^15,16^, where PI(4)P was shown to promote the formation of tubules. Thus, whether PI(4)P has a different function in the formation of tubules from autolysosomes compared to phagolysosomes remains unclear.

Recent studies have shown that vesicles carrying markers commonly found in the trans-Golgi network, including Arf1 (ADP-ribosylation factor 1), PI4KIIIβ, and TGN46, contribute to both the mitochondrial division process^17^ and the fission of Rab5-positive early endosomes^18^. Both studies showed Arf1 positive vesicles at the site of fission, and that inactivation of Arf1 or inhibition of PI4KIIIβ, which are expected to disrupt the formation of Golgi-derived vesicles, resulted in impaired mitochondria and endosome fission. Together, these studies indicate that Arf1 and PI4KIIIβ activities are required for the formation and/or function of these vesicles at the sites of fission. Based on these studies, we hypothesized that vesicles positive for Arf1 and PI4KIIIβ may also play a role in the fission of lysosomal tubules from parent lysosomal organelles. Specifically, the extensive tubulation of lysosomes observed in PI4KIIIβ-depleted cells^13^ may be due to defective fission resulting from a loss of formation and/or function of these vesicles at lysosomal tubule fission sites.

Using live-cell super-resolution microscopy, we show here that Arf1-PI4KIIIβ positive vesicles are recruited to the fission site of lysosomal tubules from several parent lysosome organelles. Our results support that they contribute to this process by mediating phosphatidylinositol-3-phosphate (PI(3)P) signaling on lysosomes at the site of fission.

## Results

### Arf1 positive vesicles and Lamp1 positive organelles form dynamic contacts

To test our hypothesis that vesicles positive for Arf1 and PI4KIIIβ contribute to the fission of lysosomal tubules, we first examined for interactions between Arf1 positive vesicles and lysosomes. We used GFP-tagged Arf1 (Arf1-GFP), which localizes to both the trans-Golgi network and vesicles and that was previously used as a marker for Arf1 positive vesicles in membrane fission events^17–19^, and the mCherry tagged late endosomal/lysosomal marker Lamp1^20,21^. Lattice light-sheet microscopy (LLSM) time-lapse imaging performed on mouse embryonic fibroblasts (MEFs) showed Lamp1 positive organelles and Arf1-GFP positive vesicles juxtaposed to each other for a prolonged time (**Fig 1a**). To determine whether these two compartments are forming membrane contact, we imaged them in live cells using a sub-Airy pinhole confocal super-resolution microscopy. We found that Arf1 positive vesicle and Lamp1 positive organelle interactions were stable and dynamic in both MEFs (**Fig 1b, c; Supplementary Movie 1**) and HeLa cells (**Sup fig 1a**). In both cell types, about 20% of Lamp1 positive organelles appeared in close proximity with at least one Arf1 positive vesicle at any given time (**Fig 1d and Sup fig 1b**) and had a mean minimum duration of contact of approximately 45s (**Fig 1e and Sup fig 1c**). Interestingly, we also observed Arf1 positive vesicles localized in juxtaposition to the site of Lamp1 positive organelle tubule fission (**Fig 1f; Supplementary Movie 2**). To explore the significance of Arf1-positive vesicles at the site of Lamp1 positive vesicle fission, we promoted the formation and fission of tubules from Lamp1 organelles by starving cells for 8h in amino acid-free media (HBSS: Hanks’ Balanced Salt Solution) as previously described^6,11^. In live cell images, we observed that Arf1-GFP positive vesicles marked more than half of the sites of tubule fission from Lamp1 positive organelles in MEFs (**Fig 1g**) and HeLa cells (**Sup fig 1d**). To ensure that the presence of Arf1 positive vesicles at Lamp1 positive tubule fission sites was not due to random positioning of the two structures, we re-quantified the same data where the Lamp1-mCherry signal was rotated by 90°. The resulting quantification was significantly lower than the original images (**Fig 1g**). Since overexpressed Lamp1 is not restricted to late endosomes/lysosomes but can also mark early endosomes^22^, we examined the cells for lysosomes tubule fission from Lamp1 positive organelles that were acidic (cresyl violet^23^ positive), or that contained overnight chased fluorescent 10kDa Dextran^24^. This revealed that Arf1-GFP positive vesicles mark lysosomal tubule fission sites (**Fig 1h and Sup fig 1e**). Taken together, our data suggest that Arf1 positive vesicles and Lamp1 organelles interact with each other and that these contacts are formed at or near the site of lysosomal tubule fission.

**Figure 1.**
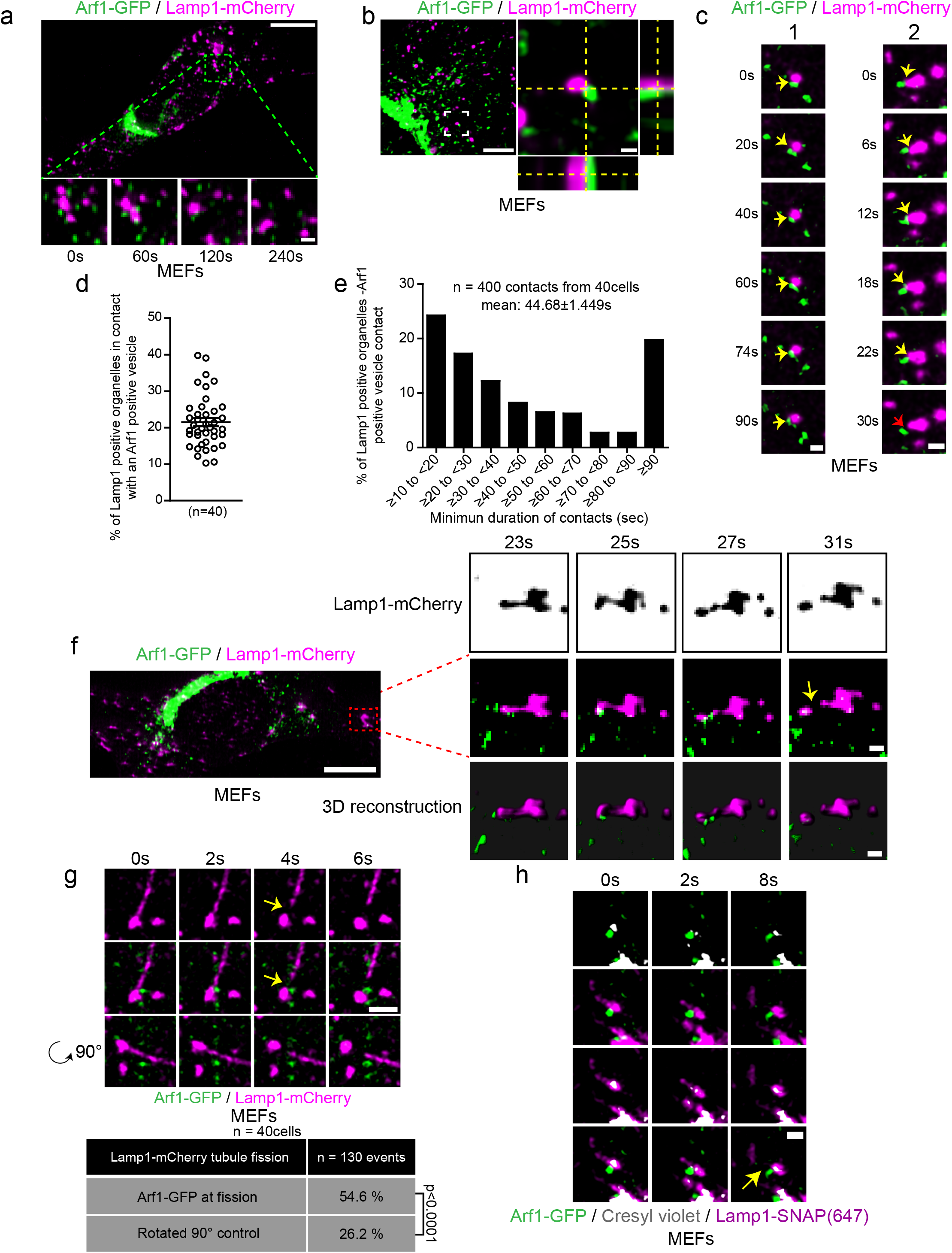
Arf1 positive vesicles make stable and dynamic contacts with Lamp1 positive organelles and mark lysosomal tubule fission sites. **a.** Lattice light sheet microscopy (LLSM) imaging of a mouse embryonic fibroblast (MEF) expressing Lamp1-mCherry and Arf1-GFP. Insets show that Arf1 positive vesicles and Lamp1 positive organelles appear to make stable contact over time. Scale bars: 10µm and 1µm (inset) **b.** Representative super-resolution image of a MEF cell expressing Arf1-GFP and Lamp1-mCherry. Inset shows a contact between an Arf1 positive vesicle and a Lamp1 positive organelle. Side panels show the view along the z-axis. Scale bar: 10µm and 1µm (inset). **c.** Representative time-lapse images of two Lamp1 positive organelle-Arf1 positive vesicle contacts. Yellow arrow indicates contact and the red arrow indicates contact untethering. Scale bars: 1µm. **d.** Quantification of the percentage of Lamp1 positive organelles in contact with at least one Arf1 positive vesicle in MEFs. The graphs show the mean ± SEM cells from three independent experiments, totaling 40 cells. **e.** Quantification of the mean minimum duration of Lamp1 positive organelle-Arf1 positive vesicle contacts in seconds. 400 randomly chosen contacts from 40 MEFs were analyzed. **f.** LLSM imaging of a MEF cell expressing Lamp1-mCherry and Arf1-GFP. Insets show time-lapse images of a single plane of only Lamp1-mCherry (inverted images) along with the corresponding merge and three-dimensional reconstruction of the same Lamp1 positive tubule fission event showing recruitment of an Arf1 positive vesicle at the fission site. Scale bars: 10µm and 1µm (insets). **g.** Representative time-lapse imaging showing an Arf1-GFP positive vesicle marking the fission site of tubule from a Lamp1-mCherry positive organelle in a MEF cell starved for 8h with HBSS (amino acid-free media). Yellow arrows indicate the fission event. The bottom panels are the same images, but the Lamp1-mCherry channel was rotated by 90°. Scale bar: 2µm. The percentage of tubule fission events marked by Arf1-GFP vesicles was quantified. n=130 events from 40 MEFs. Quantification of the same data but where Lamp1-mCherry channel was rotated by 90°. P-value from a Fisher’s exact test, two-sided unpaired t-test is shown. **h.** Representative time-lapse imaging showing an Arf1-GFP vesicle marking the fission site of a tubule from a lysosome (Lamp1-SNAP/ cresyl violet positive organelles). Yellow arrow indicates fission. Scale bar: 1µm.

### Arf1 vesicles at lysosomal tubule fission sites are positive for trans-Golgi network markers but not for endosomal markers

Since vesicles implicated in mitochondrial fission were reported to be positive for both PI4KIIIβ and TGN46^17^, we exogenously expressed fluorescent tagged versions of these proteins in MEFs and treated them with amino acid-free media (HBSS) for 8h to promote fission of tubules from Lamp1 positive organelles. We found that GFP-PI4KIIIβ positive vesicles and TGN46-mEmerald positive vesicles were recruited to approximately half of the observed Lamp1 positive organelle tubule fission events (**Fig 2a, b**), similar to that of Arf1 positive vesicles (**Fig 1g**). Importantly, we observed that vesicles recruited to tubule fission sites were positive for both Arf1 and PI4KIIIβ (**Fig 2c)**. This supports the notion that vesicles marking the sites of Lamp1 tubule fission are positive for Arf1, PI4KIIIβ and TGN46. Hereafter, we refer to these vesicles as Arf1-PI4KIIIβ positive vesicles.

**Figure 2.**
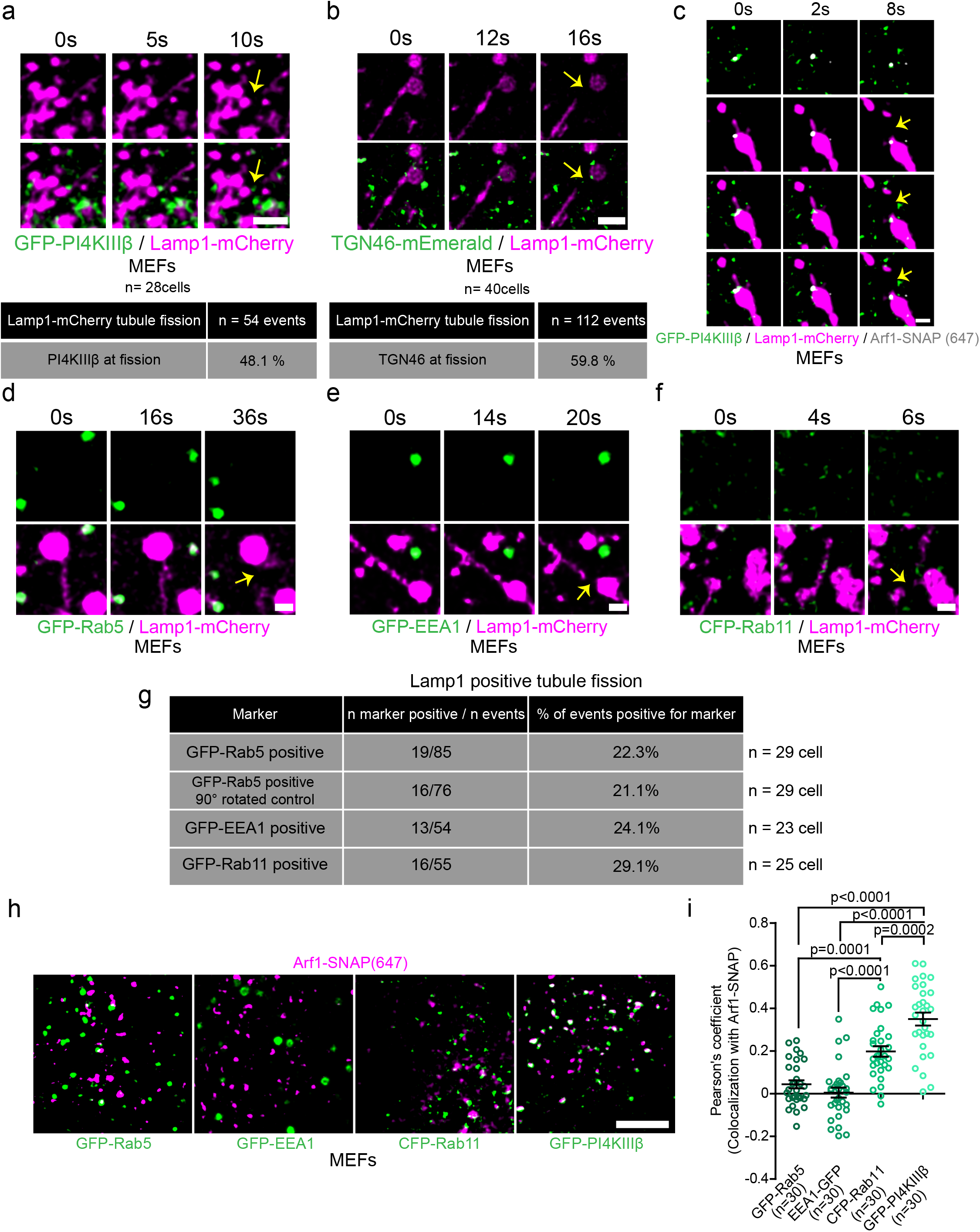
Arf1 positive vesicles recruited to Lamp1 positive organelles tubule fission sites are positive for PI4KIIIβ and TGN46 but negative for endosome markers. **a-b**. Time-lapse images of MEFs incubated for 8h in HBSS media and expressing Lamp1-mCherry and GFP-PI4KIIIβ (**a**) or TGN46-mEmerald (**b**) positive vesicles at the sites of Lamp1 positive tubule fission. Yellow arrows indicate fission. Scale bars: 2µm. Quantification of fission events with indicated vesicles (**a**) n=54 events from 28 MEFs and (**b**) n=112 events from 40 MEFs. **c**. Representative time-lapse images of a MEF cell incubated for 8h in HBSS media and expressing Lamp1-mCherry showing a Lamp1 tubules fission event marked by a vesicle positive for Arf1-SNAP and GFP-PI4KIIIβ vesicle. Yellow arrow indicates fission. Scale bar: 1µm. **d-f**. Time-lapse images of MEFs expressing Lamp1-mCherry and (**d**) GFP-Rab5, (**e**) GFP-EEA1 or (**f**) CFP-Rab11 incubated for 8h in HBSS media. Vesicles containing these proteins are absent at a Lamp1 positive tubule fission event. Yellow arrows indicate fission. Scale bars: 1µm. **g.** Quantification of such events (**d**) n=85 events from 29 MEFs, (**e**) n=76 events from 23 MEFs and (**f**) n=55 events from 25 MEFs. Also shown is the quantification of images in (**d**) where lamp1 was rotated by 90°. **h**. Representative images of a MEFs expressing Arf1-SNAP with GFP-Rab5, GFP-EEA1, CFP-Rab11 or GFP-PI4KIIIβ. Scale bar: 5µm. **i**. Pearson’s coefficient measurement of cells is described in (**h**) as indicated. The graph shows the mean ± SEM from three independent experiments. One-way ANOVA with Dunnett’s Multiple Comparison Test.

As Arf1 and TGN46 were described to localize to non-Golgi-derived vesicles such as endosomes^25,26^, we repeated Lamp1 tubule fission experiments in cells co-transfected with the early endosome markers GFP tagged Rab5 or EEA1, or with the recycling endosome marker CFP-Rab11^27^. The percentage of Lamp1 tubule fission events marked by vesicles positive for these markers was much lower than those observed for Arf1, PI4KIIIβ and TGN46 (**Fig 2d-g**). Instead, their localization to the site of fission was similar to our negative control images, where Lamp1 images were rotated by 90° (**Fig 2g**). Moreover, there was no significant colocalization of these markers with Arf1 positive vesicles in resting cells (**Fig 2h, i**), whereas we observed an extensive colocalization with PI4KIIIβ, consistent with the literature^28^. The higher colocalization between Arf1 and Rab11 compared to Rab5 suggests that some Arf1 puncta are recycling endosomes. Interestingly, Rab11 was described to localize to post-Golgi vesicles^29^, which could also explain the moderate colocalization observed here. However, the low percentage of tubule fission events marked by Rab11 compared to that of Arf1 positive vesicles indicates that the Arf1 positive vesicles recruited to fission sites are not recycling endosomes.

Taken together, our results indicate that Arf1-PI4KIIIβ positive vesicles that are recruited to Lamp1 positive organelles tubule fission sites are unlikely to be endosomal vesicles. Instead, our findings strongly suggest that they resemble more closely to vesicles derived from the Golgi Apparatus.

### Arf1-PI4KIIIβ positive vesicles mark multiple types of lysosomal tubule fission events

Lysosome reformation from autolysosomes, endolysosomes and phagolysosomes involves membrane tubulation and fission (**Fig 3a**)^1^. To address whether Arf1-PI4KIIIβ positive vesicles contribute to the fission of tubules from all of these lysosomal organelles, we first examined their recruitment at autolysosomal tubule fission sites. The formation of tubules from autolysosomes was triggered by incubating cells in amino acid-free media (HBSS) for 8 hours, at which time the formation of tubules from autolysosomes was shown to peak in multiple cell lines^6^. We identified autolysosomes as organelles positive for the membrane marker Lamp1 and the autophagic marker mCherry-LC3. We observed that Arf1 positive vesicles were present at more than half of the tubule fission events from autolysosomes in COS7 cells (**Fig 3b: Supplementary Movie 3**), suggesting that Arf1-PI4KIIIβ positive vesicles contribute to the fission of autolysosomal tubules.

**Figure 3.**
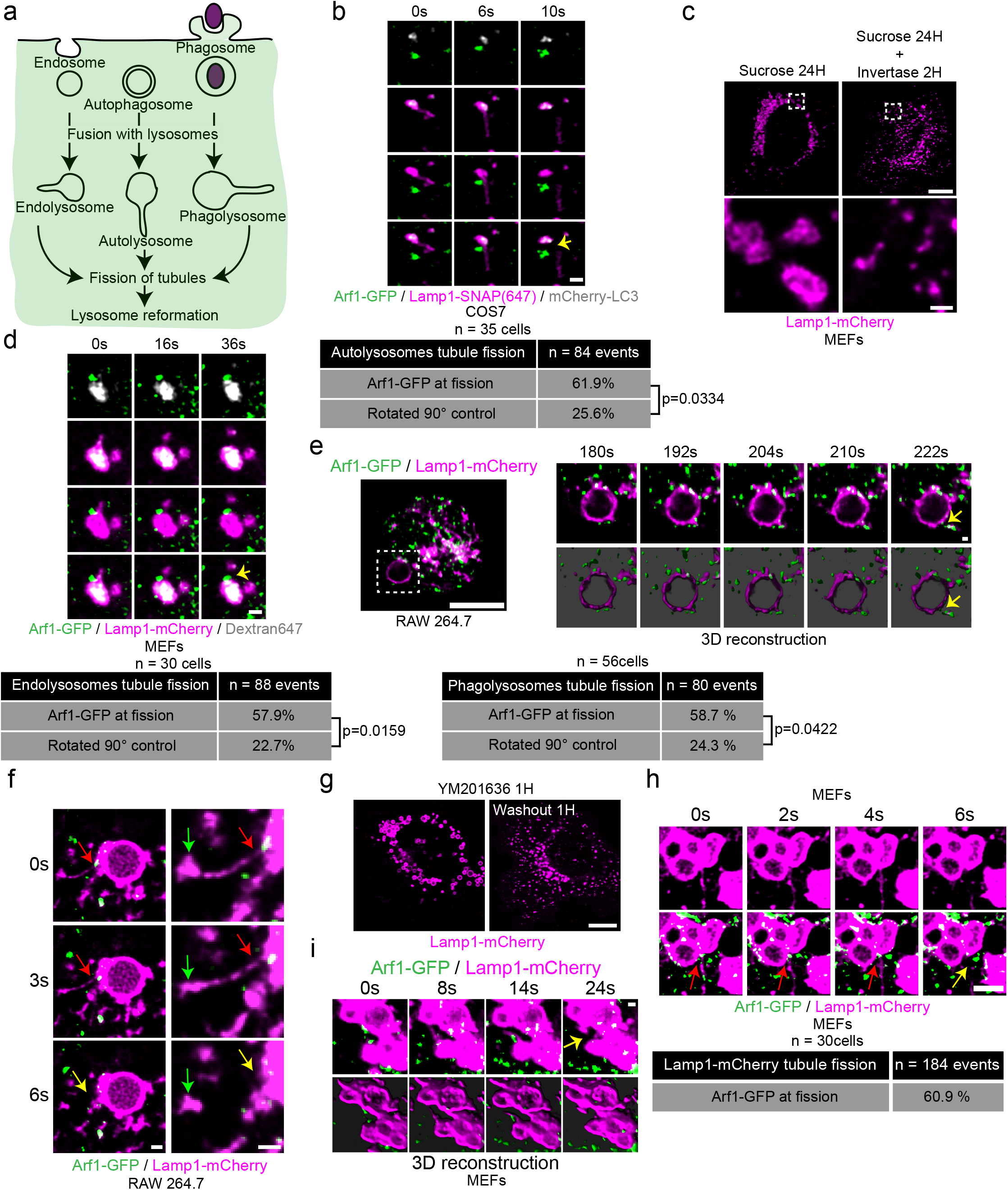
Arf1 positive vesicles mark fission sites of tubules extending from autolysosomes, endolysosomes, phagolysosomes and lysosomes. **a.** Fusion of lysosomes with endosomes, autolysosomes and phagolysosomes leads to the formation of endolysosomes, autolysosomes and phagolysosomes, respectively. Lysosomal membrane components from these lysosomal organelles are sequestered into tubule structures that detach from the organelle through a coordinated fission process to reform lysosomes. **b**. Time-lapse images of a COS7 cell showing an autolysosomal (Lamp1+/LC3+) tubule fission event marked by an Arf1 positive vesicle and quantification of such events, n=84 events from 35 COS7 cells. Yellow arrow indicates fission. Scale bar: 1µm. **c**. MEFs were incubated with 30 mM sucrose for 24 hrs to induce the formation of swollen endolysosomes (sucrosomes). Afterwards, cells were treated with Invertase (0.5mg/mL) for 2 hrs to digest sucrose and relieve endolysosome swelling and restore lysosomes. Scale bars: 10µm and 1µm (insets). **d**. Time-lapse images of MEFs showing an endolysosomal (Lamp1+/Dextran+) tubule fission event marked by an Arf1 positive vesicle and quantification of such events, n=88 events from 30 MEFs. Lamp1-mCherry signal rotated by 90° was used as a negative control. Yellow arrow indicates fission. Scale bar: 1µm. **e-f**. RAW264.7 macrophage cells fed opsonized sheep red blood cells (SRBCs), and phagocytosis synchronized via centrifugation. Live-cell timelapse images showing Arf1-GFP vesicles at phagolysosomal tubule fission sites in RAW 264.7 using LLSM and its three-dimensional reconstruction (bottom panels) (**e**) and live confocal microscopy (**f**). Red and yellow arrows indicate tubulation and tubule fission, respectively. The green arrows indicate the tip of the tubule dividing from the phagolysosome. Scale bars: 10µm for (**e**). 1µm for inset in (**e**) and (**f**). Quantification of such events, n= 80 events from 56 RAW 264.7 cells. Lamp1-mCherry signal rotated by 90° was used as a negative control. **g**. Lysosome tubule formation assay. MEFs treated with reversible PIKfyve inhibitor YM201636 (1µM for 1 hour) to promote swelling of lysosomes in a fusion-dependent manner. Washout of the inhibitor allows return to normal lysosomal size in about an hour. Scale bar: 10µm. **h-i**. Time-lapse confocal images of a Lamp1 positive tubule fission event post-YM201636 washout showing recruitment of an Arf1 positive vesicle with quantification of such events (**h**) and LLSM images with three-dimensional reconstruction (**i**). Images were acquired 5min to 40min after inhibitor washout. Red and yellow arrows indicate tubulation and tubule fission, respectively. Scale bars: 2µm (**h**) and 1µm (**i**).

As the ER was previously shown to play an important role in the fission of Rab7 positive endosomes^30^, we visualized whether it was present at the site of tubule fission marked by Arf1 positive vesicles. The ER marked almost all tubule division sites (**Sup Fig 2a**), suggesting that the recruitment and/or the function of Arf1-PI4KIIIβ positive vesicles at the fission site involves the formation of a three-way contact between the vesicles, the ER and lysosomes. As many organelles form extensive contacts with the ER^31^, it is possible that the localization of the Arf1-PI4KIIIβ positive vesicles at the sites of Lamp1 tubule fission, be a result of their co-incidence on the ER, instead of active recruitment to the site of division. To test this possibility, we monitored for the presence of peroxisomes at Lamp1 tubules fission sites as these small punctate structures appear virtually always in contact with the ER^32,33^. We found that the percentage of Lamp1 tubule fission events marked by peroxisomes (ub-RFP-SKL^34^) was not significantly different from that expected by chance (**Sup Fig 2b**). Thus, giving further support for Arf1-PI4KIIIβ positive vesicles being specifically localized to sites of tubule fission of Lamp1 positive organelles.

To examine whether Arf1-PI4KIIIβ positive vesicles also localized to fission events of tubules extruding from endolysosomes, we promoted the formation of endolysosomes by incubating cells for 24 hours in media containing 30 mM of sucrose. Sucrose is endocytosed and reaches lysosomes via the fusion of endosomes with lysosomes, forming endolysosomes called sucrosomes^7^. As lysosomes in mammalian cells are unable to digest sucrose, this disaccharide accumulates in endolysosomes inducing their swelling. The endolysosome swelling was relieved by incubating the cells with 0.5mg/mL of invertase, a yeast enzyme that hydrolyses sucrose, which reaches the swollen endolysosomes via the endocytic pathway, allowing a return to normal lysosome size (**Fig 3c**) in a reformation process implicating the formation and fission of tubules^7^. Arf1 positive vesicles were observed at more than half of tubule fission events from sucrose-induced endolysosomes (**Fig 3d**) that were identified as positive for Lamp1 and overnight chased fluorescent 10kDa Dextran (endocytosed cargo). GFP-PI4KIIIβ positive vesicles were also found at endolysosomal tubule fission sites (**Sup Fig 2c**), supporting that Arf1-PI4KIIIβ positive vesicles mark sites of tubule fission from endolysosomes.

For the recycling of phagolysosomal membrane, we used RAW 264.7 macrophages incubated with opsonized sheep red blood cells (SRBCs)^35^. We imaged RAW 264.7 cells 15 minutes after incubation with SRBCs to allow for the formation of phagolysosomes that were Lamp1 positive. Similar to the other lysosomal organelles, both Arf1-GFP and GFP-PI4KIIIβ were observed as being recruited to tubule fission events (**Fig 3e, f; Supplementary Movie 4; Sup Fig 2d**). These results support that Arf1-PI4KIIIβ positive vesicles are recruited to the fission sites of phagolysosomal tubules.

Finally, we tested whether Arf1-PI4KIIIβ positive vesicles are also recruited to fission sites of lysosomal reformation tubules without specifically promoting the formation of endolysosomes, autolysosomes or phagolysosomes. We abolished the balance between fusion and fission of lysosomes by the short-term reversible inhibition of PIKfyve, the enzyme responsible for the production of PI(3,5)P_2_ on lysosomes, using the PIKfyve inhibitor YM201636^5,36^. Inhibition of PIKfyve leads to larger and fewer lysosomes^5,36,37^. Washout of the inhibitor restored the balance between fusion and fission, leading to a recovery of lysosome size (**Fig 3g**) that involves the formation and fission of tubules. We found again that about 60% of fission events showed recruitment of an Arf1 positive vesicle (**Fig 2h, i**). These Lamp1 positive structures generated by short-term PIKfyve inhibition were negative for Rab5, indicating they are not endosome-like organelles (**Sup fig 3a**). Nor were they autolysosomes as short-term PIKfyve inhibition did not induce autophagy (**Sup fig 3b**), and the Lamp1 structures were not positive for the autophagic marker LC3 (**Sup fig 3c**). Taken together, our data show that Arf1-PI4KIIIβ positive vesicles are recruited to various types of lysosomal tubule fission events, suggesting that they contribute to the fission of lysosomal tubules.

### Arf1 inactivation and PI4KIIIβ inhibition impair the fission of lysosomal tubules

To test whether Arf1-PI4KIIIβ positive vesicles are required for the fission of lysosomal tubules, we examined whether inhibiting Arf1 activation or PI4KIIIβ function increased the number of lysosomal tubules. Such a phenotype is expected when the fission of tubules is impaired^9,12^, as demonstrated by the chemical inhibition of dynamins that contribute to tubule scission from lysosomes (**Fig 4a,b**). The chemical inhibition of PI4KIIIβ was performed with PI4KIIIbeta-IN-10^38^, and Arf1 was inactivated by treating the cells with Brefeldin A (BFA)^39^. PI4KIIIbeta-IN-10 treatment significantly reduced the number of Arf1 positive vesicles and PI4KIIIβ positive vesicles, while BFA led to a near complete loss of vesicular signal from both Arf1-GFP and GFP-PI4KIIIβ, suggesting that these treatments inhibited the formation of Arf1-PI4KIIIβ positive vesicles (**Sup Fig 4a-d**). Treatments with PI4KIIIbeta-IN-10 or BFA strongly increased the numbers of lysosomal tubules compared to vehicle control-treated cells (**Fig 4c-d**) in MEFs. Consistently, an increased number of tubules from Lamp1 positive organelles was also observed after BFA treatment in HEK293 cells (**Sup fig 4e,f**) and in primary mouse macrophages where lysosomes were marked using chased fluorescent 10 kDa Dextran (**Sup fig 4g,h**). LPS (Lipopolysaccharide) treatment, which induces striking lysosomal tubulation in primary macrophages^40^, was used as a positive control.

**Figure 4.**
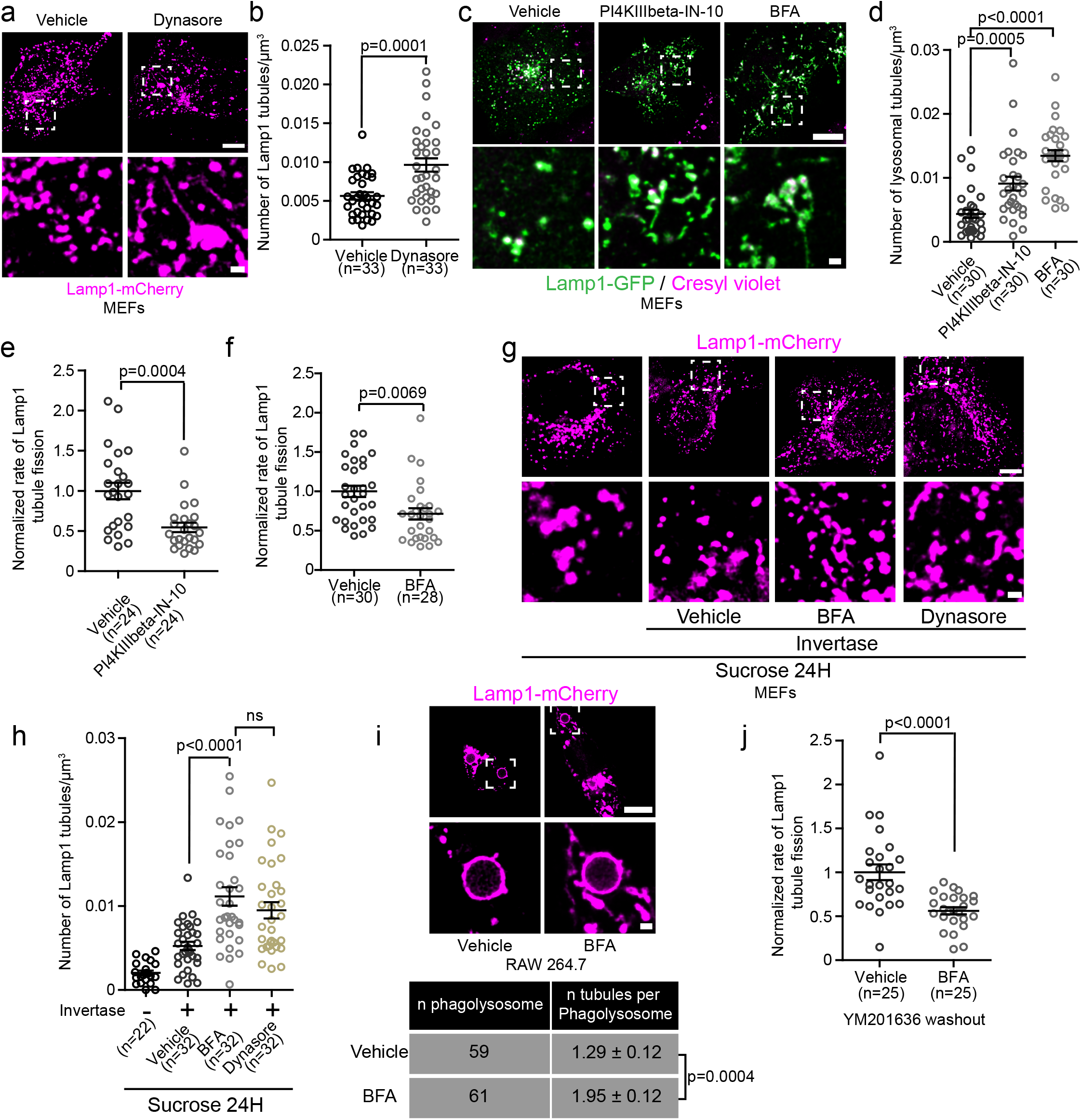
Inhibition of Arf1 activation or PI4KIIIβ function impairs the fission of lysosomal tubules. **a-b.** MEFs expressing Lamp1-mCherry treated with Dynasore (40µM) for 2 hrs show increased Lamp1 positive tubules (**a**). Scale bars: 10µm and 1µm (inset). Quantification of the number of lysosomal tubules (**b**). Two-sided unpaired t-test. **c.** MEFs treated with the PI4KIIIβ inhibitor PI4KIIIbeta-IN-10 (25nM for 3 hrs) or Arf1 activation inhibitor Brefeldin A (BFA; 10µg/mL) stained with the acidic organelle marker cresyl violet. Scale bars: 10µm and 1µm (inset). **d**. Quantification of the number of lysosomal tubules in cells described in (**c**). One-way ANOVA with Dunnett’s Multiple Comparison Test. **e-f.** Normalized rate of Lamp1 positive tubule fission in MEFs starved for 8H in HBSS treated with PI4KIIIbeta-IN-10 (25nM, 3 hrs) (**e**) or with BFA (10µg/mL, 1 hr) before imaging (**f**). Two-sided, unpaired t-test. **g**. Representative images of endolysosomes tubule formation assay in MEFs as in Fig. 3c, but after 1 hr, invertase media was replaced with media containing BFA (10µg/mL) or Dynasore (40µM) and imaged after 1 hr as indicated. Ethanol was used as vehicle control. Scale bar: 10µm. **h**. Quantification of the number of Lamp1 positive tubules of conditions in (**g**). One-way ANOVA with Dunnett’s multiple comparison test. **i**. Representative images of phagolysosomes in RAW 264.7 cells phagocyting SRBCs (scale bar: 10µm) and quantification of the number of tubules per phagosome in cells treated with BFA (10µg/mL, 30 min) or vehicle control (Ethanol, 30 min). Two-sided unpaired t-test. **j**. Normalized rate of Lamp1 positive tubule fission in MEFs after inhibition of PIKfyve (YM201636 1µM, 1 hour) and washout of the drug in the presence of BFA (10µg/mL) or vehicle control (Ethanol). Two-sided unpaired t-test. **b, d, e, f, g, h, j**. All graphs show the mean ± SEM from three independent experiments.

To determine whether the elevated number of lysosomal tubules was due to a decrease in fission, we quantified the rate of tubule fission. Both PI4KIIIbeta-IN-10 and BFA treatment significantly reduced the rate of fission (number of fission events normalized by time and cell volume) of tubules from Lamp1 positive organelles compared to vehicle-treated MEFs (**Fig 4e,f**) when autolysosomal tubules were induced by prolonged starvation (HBSS, 8 hours). Similar results were observed for endolysosomes and phagolysosomes. BFA treatment caused an increase in the number of Lamp1 tubules in sucrosomes (endolysosomes) to levels similar to cells where scission was inhibited by Dynasore treatment (**Fig 4g,h**). In RAW 264.7 cells undergoing phagocytosis of SRBCs, BFA treatment for 30min after the formation of phagolysosomes (15 minutes after phagocytosis synchronization) resulted in an increase in the number of tubules per phagolysosomes (**Fig 4i**) compared to vehicle-treated cells. We validated the change in tubules in phagolysosomes using the PI(4)P biosensor 2xP4M that was previously shown to allow visualization of tubules emerging from phagolysosomes^14^ (**Sup fig 4i**). Finally, we examined whether Arf1 was required for tubule fission in lysosomes recovering from inhibition of tubule formation. As in Fig 3, we inhibited tubule formation using the PIKfyve inhibitor YM201636. After YM201636 washout, a decrease in tubule fission rate was observed in Lamp1 positive organelles in cells treated with BFA during the washout compared to control cells (**Fig 4j**) that was accordingly linked to an increase in Lamp1 tubules (**Sup fig 4j,k**). Together, our data show that inhibiting the formation and/or function of Arf1-PI4KIIIβ positive vesicles is responsible for defects in the fission of tubules from lysosomes. Thus, giving support for a mechanism where Arf1-PI4KIIIβ positive vesicles actively contribute to the lysosomal tubule fission process.

### Arf1-PI4KIIIβ positive vesicles are not required for lysosome vesiculation or splitting

Lysosomal membrane fission occurs not only via tubulation but also by vesiculation and splitting^5^. Tubulation allows for the enrichment of membrane components and the exclusion of lumen/cargo content, while vesiculation and splitting allow for the trafficking of luminal components and cargos^5^. To test whether Arf1-PI4KIIIβ positive vesicles are also involved in fission by vesiculation and splitting, we inhibited their function and/or formation using PIK93 (PI4KIIIβ inhibitor) or BFA and followed the fragmentation of phagolysosomes containing mRFP-labelled *E. coli* in RAW 264.7 cells. After engulfment of the prey into a phagosome and fusion with lysosomes, phagolysosomes undergo extensive fragmentation that involves tubulation, vesiculation and splitting^8^. By monitoring the phagolysosome cargo (i.e. mRFP-*E. coli*) and not the membrane protein, we can specifically evaluate vesiculation and splitting forms of divisions. Interestingly, inhibition of PI4KIIIβ or Arf1 inactivation had no significant effect on phagolysosome fragmentation compared to vehicle-treated control (**Sup fig 5a,b**). Ikarugamycin, which inhibits clathrin, was used as a positive control. This indicates that Arf1-PI4KIIIβ positive vesicles are dispensable for phagolysosome fragmentation by lysosomal vesiculation and splitting. To confirm this result, we followed the fission of the Lamp1 positive organelle after PIKfyve inhibition and washout. Analysis of 345 Lamp1 positive organelles from 23 cells (15 randomly chosen Lamp1 positive organelles analyzed per cell) treated with BFA or a vehicle control showed impaired fission of tubules but not of vesicles (vesiculation) (**Sup fig 5c**).

### PI4KIIIβ inhibition increases the number of lysosomal tubules by impairing their fission

Our results suggest that inhibiting PI4KIIIβ, a factor required for the formation and/or function of Arf1-PI4KIIIβ positive vesicles, increases the number of lysosomal tubules due to impaired tubule fission (**Fig 4c,d,e**). It was previously proposed that PI4KIIIβ downregulation caused extensive lysosomal tubulation due to a loss of PI(4)P production by PI4KIIIβ at lysosomes, and thus PI(4)P produced by PI4KIIIβ at lysosomes inhibits the formation of tubules^13^. However, we find that PI(4)P was readily detected on tubules emerging from Lamp1 positive organelles after prolonged starvation and phagolysosomes (**Fig 5a,b; Sup Fig 4i**), suggesting that lysosomal tubulation caused by loss of PI4KIIIβ function could be unrelated to its potential role in PI(4)P production at lysosomes. This hypothesis is further supported by the recent report that PI4KIIIβ does not play a major role in PI(4)P synthesis at lysosomes^41^. To test whether the production of PI(4)P by PI4KIIIβ at lysosomes inhibits the formation of tubules from lysosomes, we anchored PI4KIIIβ to lysosomes by fusing it to the lysoGFP tag composed of the lysosomal anchoring sequence of p18^42^ fused to GFP (**Fig 5c**). Overexpression of LysoGFP-PI4KIIIβ in MEFs led to increased levels of the PI(4)P biosensor mCherry-P4M^43^ at Lamp1 positive organelles compared to the control Lamp1-GFP vector (**Sup Fig 6a,b**) supporting that it produced PI(4)P at lysosomes. However, lysoGFP-PI4KIIIβ was detected on Lamp1-mCherry positive tubules when formation of tubules from autolysosomes was promoted by prolonged starvation, and its mediation of an increase in lysosomal PI(4)P had no effect on the number of Lamp1-mCherry tubules compared to Lamp1-GFP expressing control cells at a basal state or after prolonged starvation (**Fig 5d,e**). These results strongly suggest that the localization of PI4KIIIβ and its production of PI(4)P at lysosomes do not inhibit the formation of autolysosomal tubules.

**Figure 5.**
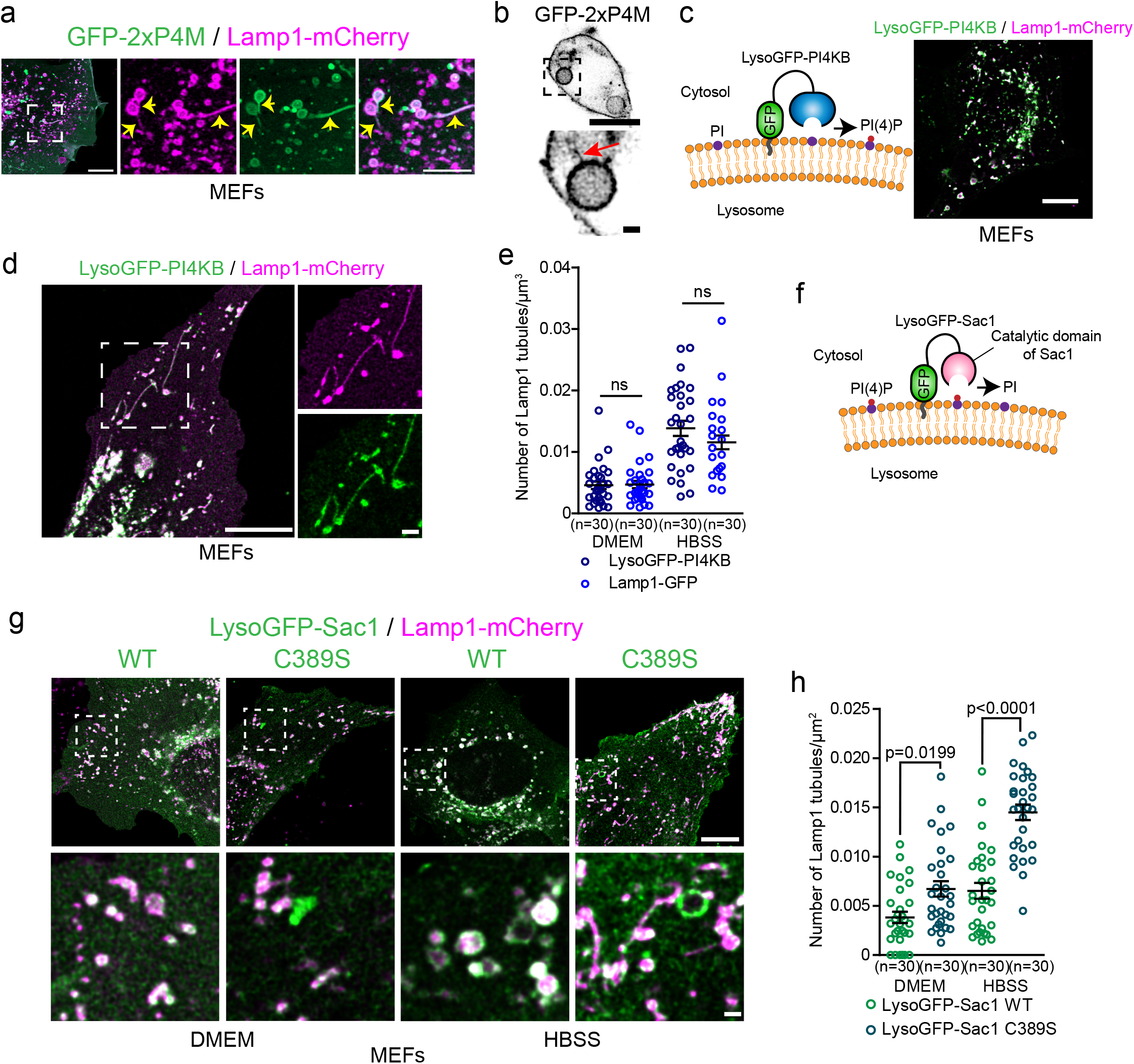
PI(4)P has a pro-tubulation role at lysosomes. **a.** Representative images of Lamp1 positive tubules in MEFs expressing Lamp1-mCherry and the PI(4)P biosensor GFP-2xP4M after 8 hrs of starvation in HBSS media. Lamp1 tubules are positive for 2xP4M. Scale bars: 10µm and 5µm (inset). **b**. Representative images of a phagolysosome showing tubules positive for the PI(4)P biosensor GFP-2xP4M in RAW 264.7 macrophages phagocyting SRBCs. Scale bars: 10 and 1µm (inset). **c**. LysoGFP-PI4KB fusion protein comprises PI(4)P producing enzyme PI4KIIIβ (PI4KB) fused to the lysosomal targeting sequence of p18. LysoGFP-PI4KB colocalizes with Lamp1-mCherry in MEFs. Scale bars: 10µm. **d**. MEFs in HBSS show LysoGFP-PI4KB colocalized with Lamp1-mCherry positive tubules. Scale bar: 10µm and 1µm (inset). **e**. Quantification of the number of Lamp1-mCherry positive tubules in MEFs expressing Lamp1-GFP or LysoGFP-PI4KB at basal state (DMEM) or 8H HBSS. The graphs show the mean ± SEM cells from three independent experiments. Two-way ANOVA with Tukey’s multiple comparison test. ns: p=0.9997 (DMEM) and p=0.3093 (HBSS). **f**. The PI(4)P phosphatase Sac1 was targeted to lysosomes by fusing it with the lysosomal targeting sequence of p18 to deplete lysosomes of PI(4)P. **g-h**. Representative images of MEFs expressing LysoGFP-Sac1 wild-type (WT) or the catalytic dead (C389S) and Lamp1-mCherry at basal state (DMEM) or after prolonged starvation (HBSS, 8H) (**g**). Scale bar: 10µm and 1µm (inset). **h**. Quantification of the number of Lamp1 positive tubules in these cells (**h**). The graphs show the mean ± SEM cells from three independent experiments. Two-way ANOVA with Tukey’s multiple comparison test.

To further clarify the role of lysosomal PI(4)P in the regulation of lysosomal tubules, we targeted the PI(4)P phosphatase Sac1 to late endosomes/lysosomes and monitored the number of Lamp1 tubules after prolonged starvation. First, we overexpressed GFP-ORPSAC1, where the catalytic domain of Sac1 replaces the lipid transfer domain of ORP1L, a late endosomal/lysosomal localized protein. This construct was previously successfully used to deplete PI(4)P levels at lysosomes^14,19^. Overexpression of ORPSAC1 but not of the catalytic dead mutant C392S led to a decreased number of Lamp1 tubules in MEFs after starvation (**Sup Fig 6c,d**). Similar but stronger inhibition was observed in both resting and starved MEF cells when lysosomal PI(4)P was depleted using the LysoGFP-Sac1 construct (**Fig 5f-h**), causing a decreased P4M signal at Lamp1 positive organelles compared to the catalytically dead mutant LysoGFP-Sac1 C389S (**Sup Fig 6e,f**). This result was also confirmed in HeLa cells (**Sup Fig 6g,h**). Finally, as PI(4)P was proposed to promote the fusion of autophagosomes with lysosomes to form autolysosomes^44^, we examined whether the expression of LysoGFP-Sac1 impaired autolysosome formation. LysoGFP-Sac1 overexpression had no effect on the colocalization of Lamp1 with the autophagosome marker mCherry-LC3 during starvation (**Sup Fig 6i,j**) or on the loss of peroxisomes by pexophagy (**Sup Fig 6k,l**), a substrate of starvation-induced autophagy^45^. These results indicate that depletion of PI(4)P by LysoGFP-Sac1 did not impair the fusion of lysosomes with autophagosomes. Thus, our data support that lysosomal PI(4)P is required to generate tubules on lysosomes and that PI4KIIIβ does not have any anti-tubulation role at lysosomes. Instead, our data suggest that PI4KIIIβ plays a role in lysosomal tubule fission by mediating the formation and/or function of Arf1-PI4KIIIβ positive vesicles that contribute to the fission of lysosomal tubules.

### The presence of Arf1-PI4KIIIβ positive vesicles at Lamp1 tubule fission sites is associated with a PI(3)P signal required for fission

Next, we investigated the role of Arf1-PI4KIIIβ positive vesicles in the lysosomal tubule fission process. Recently, a subset of Arf1 positive vesicles was shown to contribute to the fission of Rab5 positive early endosomes by promoting a phosphatidylinositol-3-phosphate (PI(3)P) increase on the dividing endosome^18^. We investigated whether Arf1-PI4KIIIβ positive vesicles could perform a similar role for the fission of lysosomal tubules. We first examined whether Lamp1 positive organelles tubule fission events were associated with a modulation of PI(3)P. We starved MEFs with amino acid-free media (HBSS) for 8h to promote the formation and fission of Lamp1 tubules and monitored PI(3)P levels on Lamp1 positive organelles undergoing tubule fission events using the PI(3)P biosensor PX-p40phox^46^ (hereafter referred to as PX). We observed that Lamp1 tubule fission events were associated with an increase in PI(3)P levels in the seconds preceding the fission event (**Fig 6a**), while there was no detectable increase on Lamp1 positive organelles with tubules that did not undergo fission (**Sup Fig 7a**). A similar increase was observed using another PI(3)P biosensor (2xFYVE), but no change was observed for a PI(4)P biosensor (2xP4M) (**Sup Fig 7b,c**). These results indicate that the increase in PI(3)P preceding fission is specific and not due to a general increase in phosphoinositides levels. The PI(3)P increase was also observed for tubule fission from lamp1 and overnight chased Dextran positive organelles, confirming that the fission of lysosomal tubule is associated with a PI(3)P increase (**Sup Fig 7d,e**).

**Figure 6.**
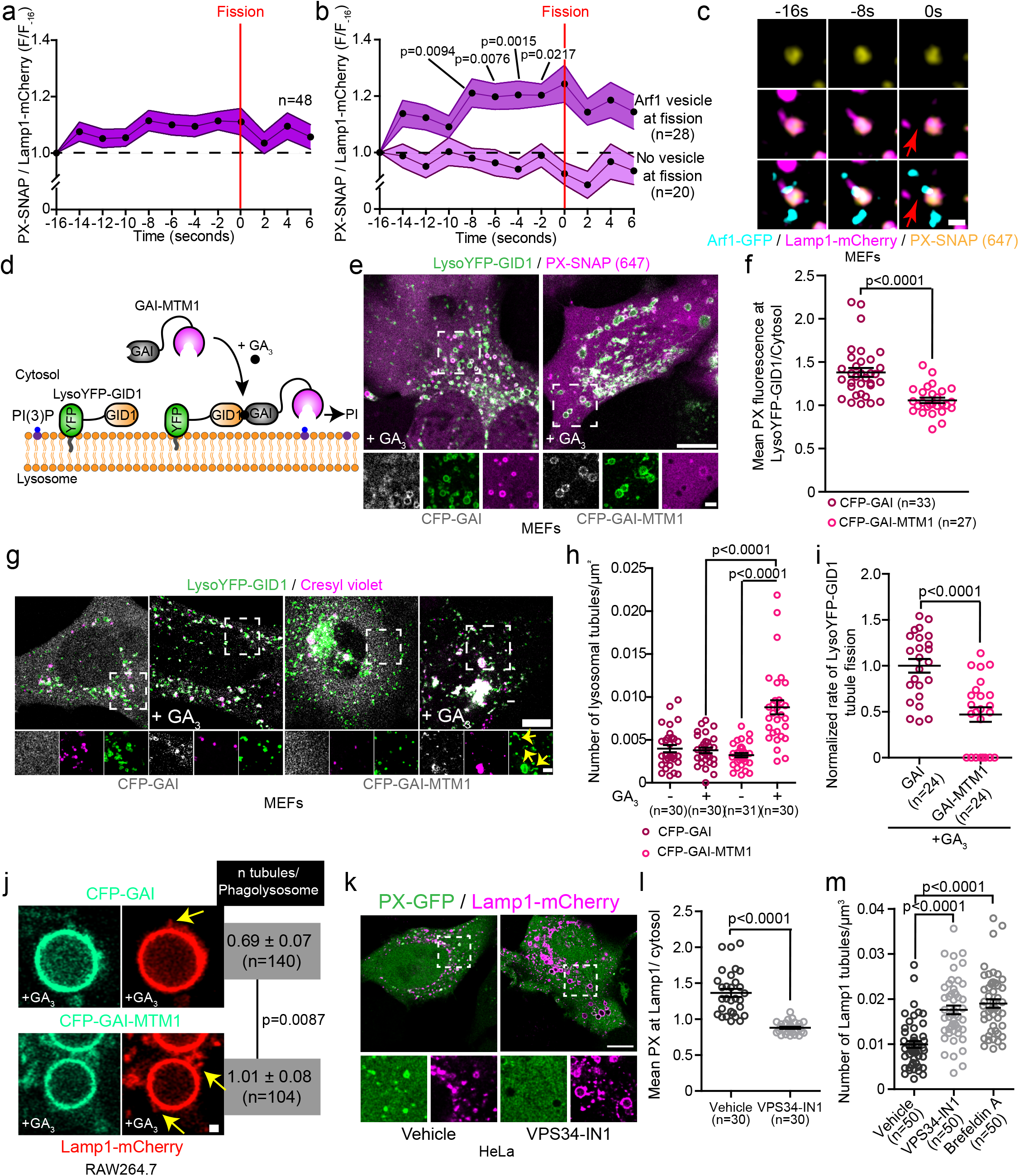
Arf1 positive vesicle-mediated Lamp1 tubules fission events are associated with a PI(3)P signal that appears required for fission. **a-c. (a)** Normalized fluorescence intensity of PX (PI(3)P biosensor) at Lamp1 positive organelles during tubule fission events. MEFs expressing Lamp1-mCherry treated with HBSS. (**b**) Events monitored in (**a**) were classified according to the presence or absence of an Arf1 positive vesicle at the fission sites. Two-way ANOVA with Tukey’s multiple comparison test. (**c**) Representative time-lapse image of a Lamp1 fission event marked by an Arf1 positive vesicles analyzed in MEFs. Red arrows indicate fission. Scale bar: 1µm. **d**. Cartoon illustration of the GAI-GID1 dimerization system used to acutely recruit the PI(3)P phosphate MTM1 to lysosomes using the LysoYFP-GID1 construct as an anchor. **e-f**. (**e**) Representative Airyscan images of MEFs expressing LysoYFP-GID1, PX-SNAP and CFP-GAI or CFP-GAI-MTM1 and treated with GA3-AM (10µM for 1H). Scale bar: 10µm and 2µm (insets) (**f**) Quantification of the mean fluorescence intensity of PX at LysoYFP-GID1 positive organelles normalized to cytosolic levels. Two-sided unpaired t-test. **g**. Representative Airyscan images of MEFs expressing LysoYFP-GID1 and CFP-GAI or CFP-GAI-MTM1 before and after treatment with GA3-AM (10µM for 1H). Acidic (Cresyl violet) LysoYFP-GID1 positive organelles were considered lysosomes. Scale bars: 10µm and 2µm (inset). Yellow arrows indicate tubules. **h**. Quantification of the number of lysosomal tubules in cells in (**g**). Two-way ANOVA with Tukey’s multiple comparison tests. **i.** Normalized rate of tubule fission from LysoYFP-GID1 positive organelles in MEFs (starved for 8h in HBSS) after recruitment of indicated construct by GA3-AM treatment (10µM) for 1 h. Two-sided unpaired t-test. **j**. Representative Airyscan images of phagolysosomes from RAW 264.7 cells phagocyting SRBCs expressing iRFP-GID1-Rab7 (not imaged), Lamp1-mCherry and CFP-GAI or CFP-GAI-MTM1 after treatment with GA3-AM (10µM for 1H) (Yellow arrows show tubules. Scale bar: 2µm) and quantification of the number of tubules per phagolysosome. Two-sided unpaired t-test. **k-m**. (**k**) Representative images of HeLa cells expressing Lamp1-mCherry and PX-GFP and treated with the VPS34 inhibitor VPS34-IN1 (1µM for 1hour) or DMSO as a vehicle control. Scale bar: 10µm and 1 µm for inset. (**l**) Quantification of the levels of PX at Lamp1 positive organelles normalized to the cytosolic level of the probe. Two-sided unpaired t-test. (**m**) Quantification of the number of Lamp1 positive tubules in these cells. BFA treatment (10µg/mL, 1H) was used as a positive control. One-way ANOVA with Dunnett’s Multiple Comparison Test. All graphs (**a**, **b, f, h, I, l, m**) show the mean ± SEM cells from three independent experiments.

Next, to determine whether Arf1-PI4KIIIβ positive vesicles are required for the observed PI(3)P increase preceding fission, we classified the Lamp1 tubule fission events in **Fig 6a** based on the presence or absence of an Arf1 positive vesicle at the site of fission. We observed that the PI(3)P increase was only observed for the fission events marked by an Arf1 positive vesicle (**Fig 6b,c**), while the other events showed no detectable PI(3)P increase. Together, these results indicate that lysosomal tubule fission events marked by Arf1-PI4KIIIβ positive vesicles are associated with a PI(3)P increase in the seconds leading to fission.

Next, we asked whether PI(3)P is required for fission by acutely depleting PI(3)P on lysosomes by targeting the PI(3)P phosphatase, MTM1, to lysosomes using the GAI-GID1 dimerization system.^47^ We generated a construct where PI(3)P phosphatase MTM1 was fused to GAI, and co-expressed it with LysoYFP-GID1 (Lyso = lysosomal targeting sequence of p18) to target it to lysosomes (**Fig 6d**). Acute recruitment of MTM1 led to a striking decrease of the PI(3)P biosensor PX levels at LysoYFP-GID1 positive organelles (**Fig 6e,f**) and to a significant increase in the number of tubules emerging from acidic LysoYFP-GID1 positive organelles (**Fig 6 g,h**) that was associated with a decrease in the rate of tubule fission from LysoYFP-GID1 positive organelles (**Fig 6i**). Similarly, the recruitment of GAI-MTM1 to phagolysosomes, using GID1-Rab7 as an anchor^14^, resulted in an increased number of tubules per phagolysosomes in RAW264.7 macrophages (**Fig 6j**). To confirm the importance of PI(3)P in the fission of lysosomal tubules, we inhibited the production of PI(3)P by VPS34, a PI3-kinase that notably produces PI(3)P at lysosomes^48^. This led to a strong depletion of PI(3)P levels at Lamp1 positive organelles and markedly increased the number of Lamp1 tubules in HeLa cells (**Fig 6k-m**), as previously reported^48^, while it did not affect the number of Arf1 or PI4KIIIβ positive vesicles (**Sup Fig 7f-i**). These results support that PI(3)P is required on lysosomes for tubule fission. Collectively, these data suggest that Arf1-PI4KIIIβ positive vesicles at the lysosomal tubule fission events mediate the increase in PI(3)P on the lysosome that drives fission.

### The Arf1-PI4KIIIβ positive vesicle localized protein SEC14L2 contributes to the fission of lysosomal tubules

The PI(3)P signalling involved in Rab5 early endosomes fission regulated by Arf1 positive vesicles was shown to be mediated by a protein called SEC14L2^18^. SEC14L2, which is found on Arf1 positive vesicles, is a lipid-binding protein that binds and transfers PI(3)P. SEC14L2 was proposed to regulate Rab5 early endosomes fission by either transferring PI(3)P from Arf1 vesicles to Rab5 early endosomes or by activating the PI(3)P production machinery at the endosome-Arf1 vesicle contact site^18^. Thus, we asked whether SEC14L2 could also contribute to the fission of lysosomal tubules mediated by Arf1-PI4KIIIβ positive vesicles. When overexpressed, SEC14L2 shows a diffuse cytosolic signal in MEFs. However, the removal of excess cytosolic signal by permeabilizing the plasma membrane with digitonin before fixation showed colocalization of SEC14L2-mCherry to a subpopulation of Arf1-PI4KIIIβ positive vesicles in HeLa cells (**Fig 7a**) suggesting that SEC14L2 localizes to a subset of Arf1-PI4KIIIβ positive vesicles.

**Figure 7.**
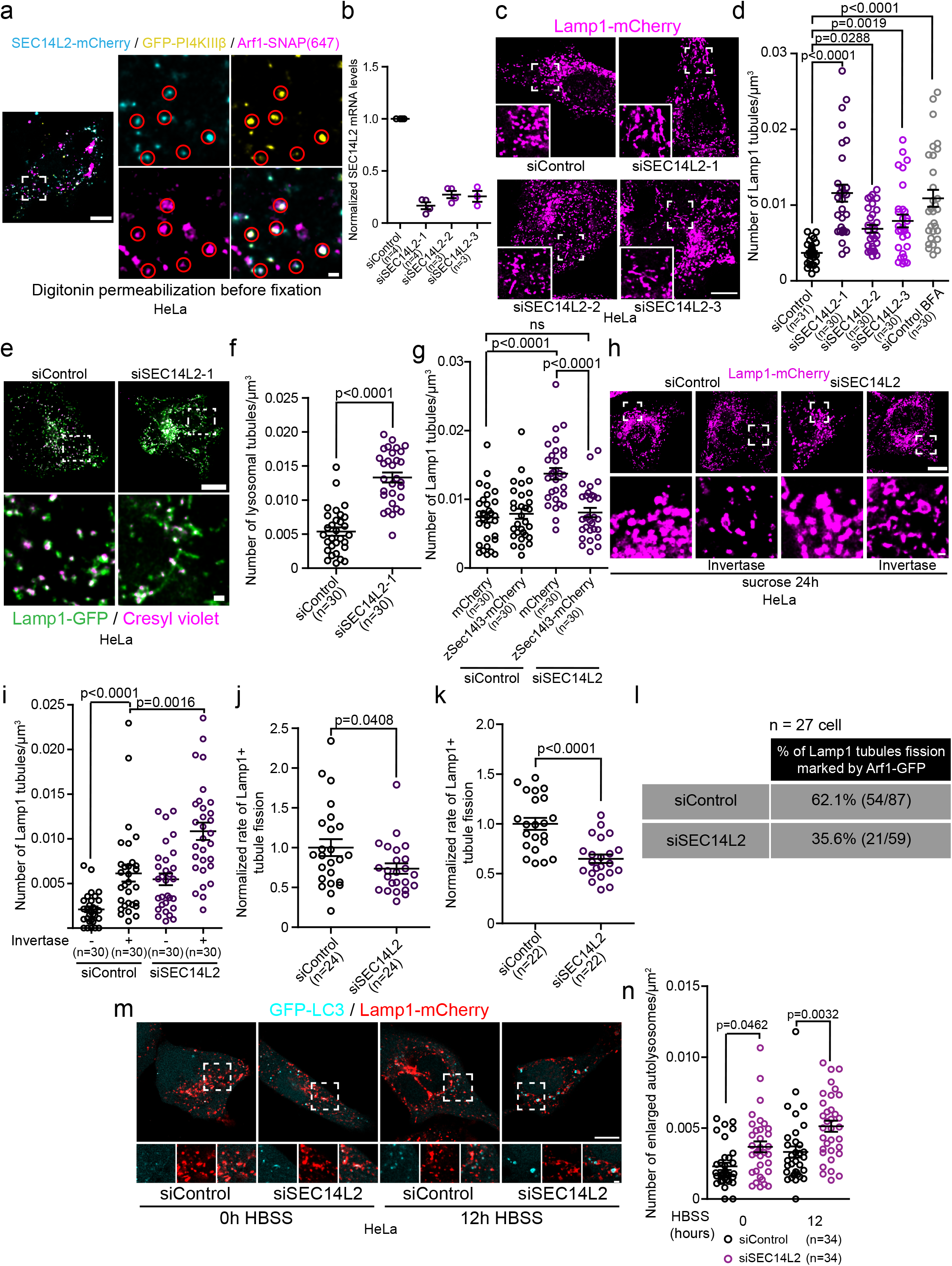
The Arf1-PI4KIIIβ positive vesicle localized protein SEC14L2 contributes to the fission of lysosomal tubules. **a.** Representative live image of a HeLa cell expressing SEC14L2-mCherry, GFP-PI4KIIIβ and Arf1-SNAP. Excess cytosolic signal was removed by short-term permeabilization with digitonin before fixation. Red circles show SEC14L2 colocalization with Arf1-PI4KIIIβ positive vesicles. Scale bar: 10µm and 1µm (inset). **b**. Normalized SEC14L2 mRNA levels in HeLa cells treated with the indicated siRNAs. **c-d**. Representative images of Lamp1 positive organelles morphology in HeLa cells expressing Lamp1-mCherry and treated with the indicated siRNAs (**c**) and quantification of the number of Lamp1 positive tubules in these cells (**d**). Scale bar: 10µm. BFA treatment (10µg/mL for 1 hour) was used as a positive control. One-way ANOVA with Dunnett’s Multiple Comparison Test. **e-f**. Representative images of lysosomes (Cresyl violet positive Lamp1 organelles) in HeLa cells treated with the indicated siRNAs (**e**) and quantification of the number of lysosomal tubules (**f**). Scale bar: 10µm and 1µm (inset). Two-sided unpaired t-test. **g**. Quantification of the number of Lamp1 positive tubules in HeLa cells expressing cytosolic mCherry or the zebrafish SEC14L2 homologue zSec14l3-mCherry after treatment with the indicated siRNAs. Two-way ANOVA with Tukey’s multiple comparisons test. ns: p-value=0.8998. **h**. Representative images of HeLa cells treated with the indicated siRNAs and incubated with 30mM sucrose for 24h to promote the formation of endolysosomes (sucrosomes) before and after 2 hrs Invertase (0.5mg/mL) as indicated. Scale bars: 10µm and 1µm (inset). **i.** Quantification of the number of Lamp1 positive tubules in these cells in (**h**). Two-way ANOVA with Tukey’s multiple comparisons test. **j-k**. Normalized rate of Lamp1 tubule fission in HeLa cells treated with the indicated siRNAs after prolonged starvation (HBSS 8H) (**j**) or short-term inhibition of PIKfyve using 1µM of YM201636 and washout of the drug (**k**). Two-sided unpaired t-test. **l**. Quantification of the number of Lamp1 positive tubule fission events marked by an Arf1 positive vesicle in HeLa cells treated with indicated siRNAs and starved (HBSS) for 8 hours. **m.** Representative images of HeLa cells expressing CFP-LC3 and Lamp1-mCherry treated with the indicated siRNAs and incubated in HBSS for the indicated time. Scale bars: 10µm and 1µm. **n**. Quantification of the number of enlarged autolysosomes (>1µm²) of cells in (**m**). Two-Way ANOVA with Tukey’s multiple comparison tests. All graphs (**b, d, f, g, i, j, k, n**) show the mean ± SEM from three independent experiments.

To assess whether SEC14L2 was required for lysosome tubule fission, we depleted the cells of endogenous SEC14L2 using siRNAs (**Fig 7b**). We found that depleting cells of SEC14L2 significantly increased the number of tubules from Lamp1 positive organelles compared to siRNA control-treated cells in HeLa cells and at similar levels to BFA treatment (**Fig 7c,d**). Cresyl violet staining validated that increased Lamp1 tubules were lysosomal (**Fig 7e,f**). To check for potential off-target effects of the siRNA, we expressed the zebrafish homologue of human SEC14L2, zSec14l3^18^, and found that it rescued the increase in tubules induced by the knockdown (**Fig 7g**). A similar increase in tubulation caused by SEC14L2 knockdown was observed for invertase-treated sucrosomes (endolysosomes) (**Fig 7h, i**).

To confirm that the increased number of lysosomal tubules in SEC14L2 depleted cells was caused by defective fission, we monitored the rate of tubules fission from Lamp1 positive organelles fission induced by prolonged starvation or YM201636 treatment and washout. In both cases, siRNA-mediated depletion of SEC14L2 caused a significant decrease in the rate of tubule fission (**Fig 7j,k**). Moreover, we observed that the percentage of Lamp1 tubule fission event marked by an Arf1 positive vesicle was reduced in HeLa cells depleted of SEC14L2 compared to cells treated with a control siRNA (**Fig 7l**). Lastly, the depletion of SEC14L2 depletion did not affect the formation of Arf1 or PI4KIIIβ positive vesicles (**Sup fig 7j-m**), suggesting that the decrease in fission events is not due to a loss of the Arf1-PI4KIIIβ vesicles. Taken together, our results support that SEC14L2, which localizes on Arf1-PI4KIIIβ positive vesicles, is required for the fission of lysosomal tubules.

Finally, since the formation and fission of autolysosomal tubules are required for clearance of enlarged autolysosomes after prolonged starvation (typically 12h in HBSS)^11^, a defect in tubule fission would prevent the clearance of these enlarged autolysosomes. Thus, to confirm a role for SEC14L2 in the fission of lysosomal tubules, we monitored the number of enlarged autolysosomes in HeLa cells depleted of SEC14L2. We found that HeLa cells depleted of SEC14L2 had significantly more enlarged autolysosomes after prolonged starvation compared to siRNA control-treated cells (**Fig 7m,n)**. This result indicates that the knockdown of SEC14L2 impairs autophagic lysosome reformation and further supports that SEC14L2 is required fission of lysosomal tubules.

### The PI(3)P binding domain of SEC14L2 is required for regulating lysosomal PI(3)P for lysosomal tubule fission

A key functional domain of SEC14L2 is its lipid binding CRAL-TRIO domain, which is required for binding and transferring PI(3)P^18^. To test whether SEC14L2’s lipid binding/transfer activity is required for lysosomal tubule fission, we performed complementation experiments by reintroducing various deleted constructs of zSec14l3 to HeLa cells depleted of its endogenous SEC14L2 with siRNA. Three different zSec14l3 mutants were used: (i) the previously described PI(3)P binding deficient mutant M5^18^; (ii) the ΔCRAL-TRIO construct that lacks the lipid binding domain, and (iii) the ΔGOLD construct lacking the Golgi dynamics (GOLD) domain of the protein (**Fig 8a**). We found that only the overexpression of wild-type zSec14l3 decreased the number of Lamp1 positive organelles tubules in cells depleted of SEC14L2 (**Fig 8b**). This strongly suggests that both its PI(3)P binding ability and GOLD domain are crucial for the fission of lysosomal tubules. Accordingly, the rate of tubule fission from Lamp1 positive organelles in HeLa cells depleted of SEC14L2, which is reduced compared to control cells (**Fig 7j**), was increased by the expression of WT zSec14l3 but not by that of the PI(3)P binding deficient mutant M5 (**Fig 8c**).

**Figure 8.**
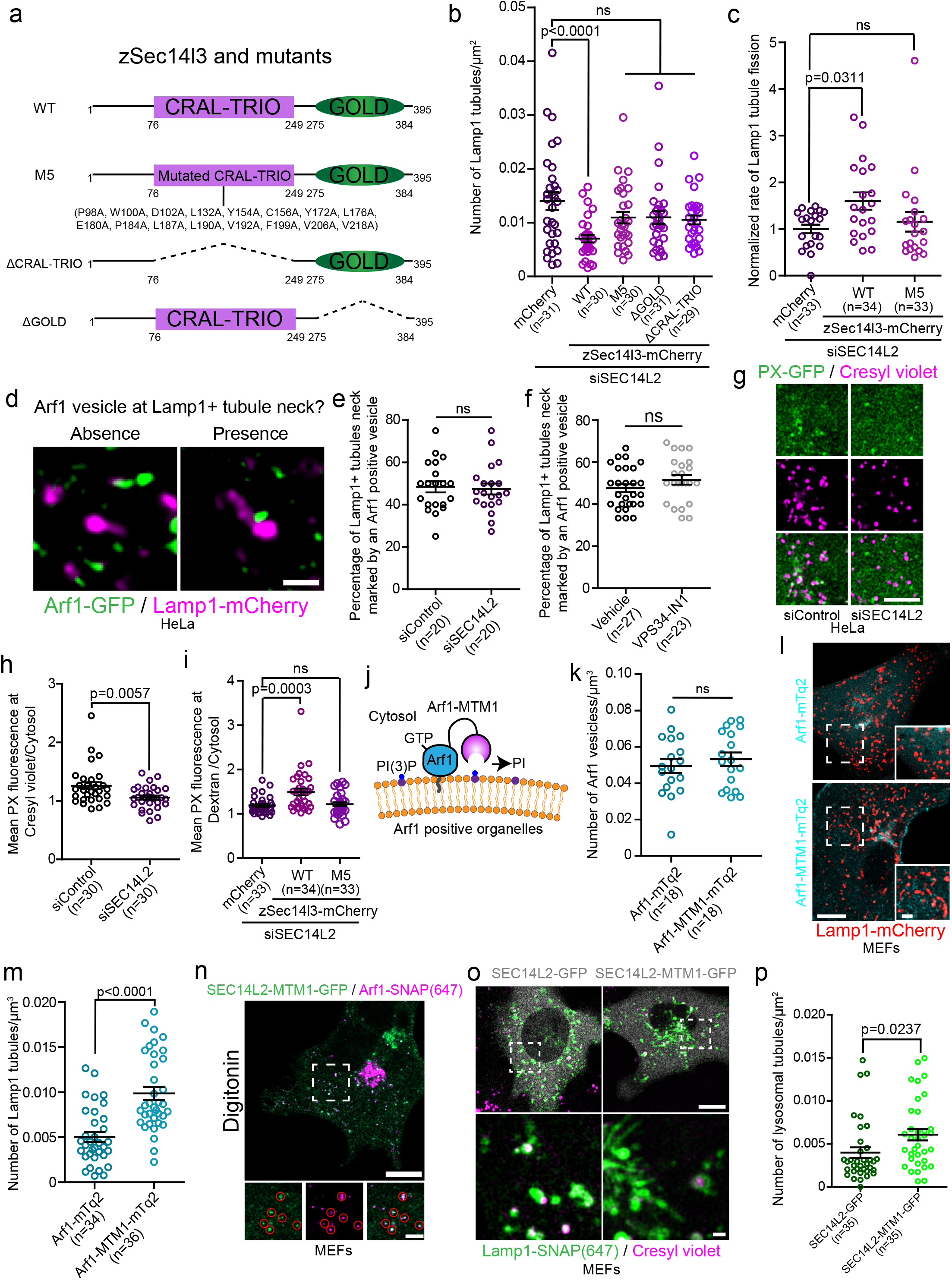
The PI(3)P binding domain of SEC14L2 is required for regulating lysosomal PI(3)P and tubule fission. **a.** Cartoon illustration of the various zSec14l3 constructs used in the study. **b.** Quantification of the number of Lamp1 positive tubules in cells treated with indicated siRNA and expressing the indicated constructs. One-way ANOVA with Dunnett’s Multiple Comparison Test. p-value = 0.1776 (M5), 0.1808 (ΔCRAL-TRIO) and 0.1069 (ΔGOLD). **c**. Normalized rate of Lamp1 positive tubule fission in cells treated with indicated siRNA and expressing the indicated constructs. One-way ANOVA with Dunnett’s Multiple Comparison Test. ns = 0.7442 **d-f**. Representative images showing the absence and presence of an Arf1 positive vesicle at Lamp1 positive organelle tubule necks. Scale bar: 1µm (**d**). Quantification of the percentage of Lamp1 tubules necks marked by an Arf1 positive vesicle in HeLa cells treated with the indicated siRNA and starved (HBSS) for 8 hours (**e**) or treated with VPS34-IN1 (1µM for 1 hour) (**f**). Two-sided unpaired t-test. ns: p-value = 0.7773(**e**) and 0.1825 (**f**). **g.** Representative images of a 10 µm × 10 µm section of a HeLa cell treated with the indicated siRNA expressing the PI(3)P biosensor PX-GFP and where acidic lysosomes were marked using cresyl violet Scale bar: 5µm. **h.** Quantification of the PX levels colocalizing with cresyl violet normalized to the cytosolic level of the probe of cells in (**g**). Two-sided unpaired t-test. **i**. Quantification of the PX levels colocalizing with overnight chased 10kDa fluorescent Dextran normalized to the cytosolic level of the probe in HeLa cells treated with indicated siRNA and expressing the indicated constructs. One-way ANOVA with Dunnett’s Multiple Comparison Test. ns = 0.8898. **j**. The PI(3)P phosphatase MTM1 can be anchored at Arf1 positive organelles by fusing it to Arf1, allowing the depletion of PI(3)P at these structures. **k**. Quantification of the number of Arf1-GFP positive vesicles per cell in MEFs expressing Arf1-mTq2 or Arf1-MTM1-mTq2. Two-sided unpaired t-test. ns: p-value = 0.4901. **l-m**. (**l**) Representative images of s expressing Lamp1-mCherry and Arf1-mTq2 or Arf1-MTM1-mTq2, scale bar: 10µm, and (**m**) quantification of the number of Lamp1 positive tubules in these cells. Two-sided unpaired t-test. **n**. Representative Airyscan image of a MEF cell expressing SEC14L2-MTM1-GFP and Arf1-SNAP. Red circles show SEC14L2-MTM1 colocalizing with Arf1 positive vesicles. Scale bar: 10µm and 1µm (inset). **o-p**. Representative Airyscan images of MEFs expressing Lamp1-SNAP and SEC14L2-GFP or SEC14L2-MTM1-GFP and treated with Cresyl violet to mark acidic lysosomes (scale bar: 10µm and 1µm (inset)) (**o**) and the quantification of the number of lysosomal tubules (**p**). Two-sided unpaired t-test. All graphs (**b, c, e, f, h, i, k, m, p**) show the mean ± SEM cells from three independent experiments.

Since we found SEC14L2 localized on Arf1-PI4KIIIβ positive vesicles and its PI(3)P binding domain was required for the efficient fission of tubules from Lamp1 positive organelles, a possible function for SEC14L2 during tubule fission may be to mediate the recruitment of Arf1-PI4KIIIβ positive vesicles to the site of lysosomal tubule fission by binding to lysosomal PI(3)P. To test this possibility, we monitored for the presence of Arf1 positive vesicles at Lamp1 positive organelles tubule necks, where we observed the vast majority of fission events. We found that there was no difference in the percentage of Lamp1-positive organelle tubule necks showing contact with an Arf1 positive vesicle in SEC14L2-depleted cells compared to the control siRNA-treated cells (**Fig 8d-e**). This suggests that SEC14L2 has no role in the recruitment of Arf1-PI4KIIIβ positive vesicles to the site of lysosomal tubule fission. Similar results were observed in cells treated with the VPS34 inhibitor (**Fig 8f**), suggesting that an increase in lysosomal PI(3)P is not necessary for Arf1-PI4KIIIβ positive vesicle recruitment to the fission.

Since we find that the PI(3)P binding of SEC14L2 is required for Lamp1 positive tubule fission and that lysosomal tubules fission events marked by an Arf1 positive vesicle are associated with an up-regulation of lysosomal PI(3)P levels, we reasoned that SEC14L2 could mediate the PI(3)P signalling required for tubule fission. Consistent with this hypothesis, depletion of SEC14L2 led to a decrease in the PI(3)P biosensor PX fluorescence at acidic lysosomes in resting cells, compared to cells treated with a control siRNA (**Fig 8g,h**). Importantly, the decrease in PI(3)P at lysosomes was rescued by the overexpression of wild-type zSec14l3 but not of the PI(3)P binding deficient mutant M5 in cells where lysosomes were identified using overnight chased 10kDa Dextran (**Fig 8i**). These results support that SEC14L2 contributes to the fission of lysosomal tubules by regulating the levels of lysosomal PI(3)P.

Finally, SEC14L2 was shown to transfer PI(3)P and to promote PI(3)P production by activating VPS34 *in vitro*^18^. Thus, we reasoned that if SEC14L2 regulates lysosomal PI(3)P to promote tubule fission by transferring it from Arf1-PI4KIIIβ positive vesicles or by activating VPS34 on lysosomes, the depletion of PI(3)P in the immediate proximity of SEC14L2 should impair tubule fission and leads to an increase in the number of Lamp1 positive tubules. To test this, we first fused the PI(3)P phosphatase MTM1 to Arf1 (**Fig 8j**). Anchoring MTM1 to Arf1 did not affect the formation of Arf1 positive vesicles (**Fig 8k**) but led to an increased number of Lamp1 positive tubules in MEFs as compared to cells expressing Arf1-mTq2 that were used as a control (**Fig 8l,m**). Second, we fused MTM1 directly to SEC14L2, which did not impair the localization of SEC14L2 to Arf1 positive vesicles (**Fig 8n**), but did lead to an increase in the number of lysosomal tubules (**Fig 8o,p**). These results strengthen the link between SEC14L2 and PI(3)P in lysosomal tubule fission events as they support that depleting PI(3)P at the site of SEC14L2 localization or its immediate proximity impairs lysosomal tubule fission. Further, this result is consistent with a model where SEC14L2 regulates lysosomal PI(3)P levels to promote tubule fission by a transfer mechanism from Arf1-PI4KIIIβ positive vesicles or by activating VPS34 on lysosomes at contact sites with the vesicles. Collectively, our data suggest that Arf1-PI4KIIIβ positive vesicle localized SEC14L2 is required for the fission of lysosomal tubules by mediating a PI(3)P signalling on lysosomes.

## Discussion

Formation and fission of lysosomal tubules allow for the reformation of lysosomes from parent lysosome organelles such as endolysosomes, autolysosomes or phagolysosomes^1,6–8^. However, the molecular mechanisms regulating these processes are only partially known. In this study, we used super-resolution live-cell imaging to investigate the role of Arf1-PI4KIIIβ positive vesicles in lysosomal tubule fission events. We showed that these vesicles are recruited to a broad range of lysosomal tubule fission events and that inactivation of Arf1 or inhibition of PI4KIIIβ impairs the fission of lysosomal tubules. Our study also suggests that Arf1-PI4KIIIβ positive vesicles mediate a PI(3)P signal at the site of fission through a SEC14L2-dependent mechanism. This mechanism involves SEC14L2 either mediating a transfer of PI(3)P from the vesicle to the lysosome or activating lysosomal VPS34 at contact sites between the vesicle and the lysosome. **Figure 9** provides a visual representation of this proposed mechanism.

**Figure 9.**
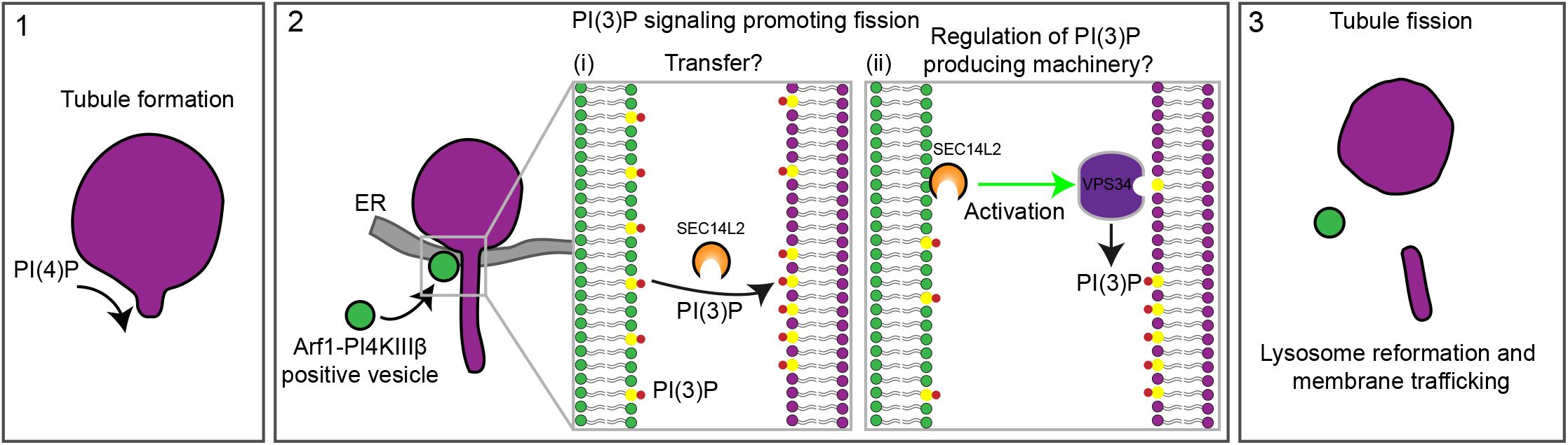
Proposed model for Arf1-PI4KIIIβ positive vesicles function at lysosomal tubules fission site. **(1)** Formation of lysosomal tubules from lysosomes, autolysosomes, endolysosomes, and phagolysosomes requires PI(4)P on the cytosolic leaflet of the lysosome membrane. **(2)** Arf1-PI4KIIIβ positive vesicles are recruited to lysosomal tubule fission events, where the ER is also present. At these tri-organelle contact sites, we propose that SEC14L2 on the Arf1-PI4KIIIβ positive vesicle mediates a PI(3)P signal on the lysosomal membrane by (i) transferring PI(3)P from the vesicle to the lysosome, or (ii) regulating the activity of PI(3)P production machinery on lysosomes. These two mechanisms are not mutually exclusive. **(3)** PI(3)P signaling on tubules leads to fission of the tubule allowing for lysosome reformation from parent lysosome organelles (autolysosomes, endolysosomes or phagolysosomes) or membrane trafficking events. Lysosomes are depicted in magenta, Arf1-PI4KIIIβ positive vesicles in green and the ER in gray.

The vesicles that mark the site of lysosomal tubule fission are positive for several markers associated with the Golgi apparatus, Arf1, PI4KIIIβ and TGN46, suggesting that these vesicles likely derive from the Golgi Apparatus. Similar Golgi protein positive vesicles were previously reported in the fission of endosomes^18^ and mitochondria^17^ and were termed Golgi-derived vesicles. However, we elected to call them Arf1-PI4KIIIβ positive vesicles since none of the markers used in our study are restricted to the Golgi Apparatus and Golgi-derived vesicles. For instance, both Arf1 and TGN46 have been shown on endosomes, albeit at low amounts^25,26^, thus it is possible that these vesicles at the site of lysosomal tubule fission could represent non-Golgi pools of Arf1, PI4KIIIβ and TGN46. However, they were negative for several endosomal markers suggesting these Arf1-PI4KIIIβ positive vesicles are not of endosomal nature. Further studies are required to determine the precise nature of these vesicles, including whether they represent Golgi-derived vesicles and whether they differ from the Arf1 positive vesicles regulating the fission of endosomes or mitochondria.

The recruitment of Arf1-PI4KIIIβ positive vesicles to fission sites likely involves various tethering factors allowing the establishment of a membrane contact site between the vesicle, the lysosome and potentially the ER at the site of fission. Indeed, we observed that the ER marked the majority of Lamp1 tubule fission sites suggesting that a three-way contact between the three structures is involved. Multi-organelle interactions were recently identified in the division of mitochondria^17,19^ and endosomes^18^, further suggesting that membrane fission events are highly coordinated processes requiring the involvement of multiple organelles. In all these cases, the ER appears to be the common organelle, thus supporting the idea that the ER may regulate contact between different organelles^19,49^.

Lysosomes undergo several different forms of fission events, including tubulation, vesiculation and splitting^5–8,13^. We found that Arf1-PI4KIIIβ positive vesicles contribute only to the fission of tubules and are dispensable for the fission of vesicles or by splitting of lysosomes, suggesting a mechanistic, regulatory and functional difference between tubule fission versus the other division pathways. The main difference is likely due to the need to recycle membranes and membrane proteins without contamination of the lysosomal lumen content. However, further studies would be required to investigate the different mechanisms in play for the various types of lysosomal fission events.

Interestingly, we observed that Arf1-PI4KIIIβ positive vesicles are recruited to approximately 60% of the tubule fission events in autolysosomes, phagolysosomes and endolysosomes. This implies that Arf1-PI4KIIIβ positive vesicles could act as a facilitator for the membrane scission event or that several types of mechanistically and functionally distinct lysosomal tubule fission events may exist, similar to what was recently reported for mitochondria^50^. Consistent with this hypothesis, while lysosomal tubulation and fission were most often described in the process of lysosome reformation^6–8^, such events may also allow for the traffic of lipids such as cholesterol^51^.

One possible reason for the recruitment of Arf1-PI4KIIIβ positive vesicles may be to regulate the PI(3)P increase at this fission site. A recent study reported that Arf1 positive Golgi-derived vesicles contribute to the division of endosomes by promoting an increase of PI(3)P on their membranes. We found that lysosomal tubule fission events marked by Arf1-PI4KIIIβ positive vesicles were similarly associated with an increase in PI(3)P levels on lysosomes in the seconds preceding fission. The increase in PI(3)P is not required for recruiting Arf1-PI4KIIIβ positive vesicles to the lysosomal tubule fission site since decreasing lysosomal PI(3)P levels by inhibiting VPS34 did not alter the recruitment of Arf1-PI4KIIIβ positive vesicles to Lamp1 positive tubule necks. Moreover, acute depletion of PI(3)P at organelles positive for the lyso tag (P18 lysosomal anchoring n-terminus)^42^ markedly increased the number of lysosomal tubules in cells, suggesting a critical role for PI(3)P in the process of tubule fission. Collectively, these data support that Arf1-PI4KIIIβ positive vesicles mediate a PI(3)P signalling to drive the fission of lysosomal tubules.

We identified the PI(3)P binding protein SEC14L2 as a potential mediator of the increase of PI(3)P at the site of fission. SEC14L2 is a lipid-binding protein that was shown to bind and transfer PI(3)P *in vitro*^18^. Further, Arf1 positive vesicle localized SEC14L2 was shown to contribute to the fission of Rab5 early endosomes via the regulation of endosomal PI(3)P^18^. We find that SEC14L2 localizes to a subset of Arf1-PI4KIIIβ positive vesicles and is required for efficient fission of lysosomal tubules, as depleting SEC14L2 impaired the fission of tubules. Further, the depletion of SEC14L2 decreased PI(3)P levels on lysosomes. Overexpression of wild-type zSec14l3, a zebrafish homologue of SEC14L2, but not of a PI(3)P binding deficient mutant in cells depleted of SEC14L2 increased both the Lamp1 tubule fission rate and lysosomal PI(3)P levels. These findings support a model where the Arf1-PI4KIIIβ positive vesicle localized SEC14L2 regulate lysosomal PI(3)P to promote the fission of their tubules.

There are several models in which SEC14L2 can regulate PI(3)P at the site of lysosome tubule fission. One possible model for SEC14L2 function at the site of fission is to provide PI(3)P to lysosomes by transferring it from Arf1-PI4KIIIβ positive vesicles. This is supported by recent reports that SEC14L2 can transport PI(3)P between liposomes^18^ and our findings that fusing the PI(3)P phosphatase MTM1 to SEC14L2 or Arf1 to deplete PI(3)P in the immediate proximity to SEC14L2 increases the number of Lamp1 positive tubules. However, it is equally possible that SEC14L2 increases PI(3)P on lysosomes by regulating the production of PI(3)P on lysosomes in a mechanism dependent on its ability to bind PI(3)P. In favor of this model, SEC14L2 was shown to activate VPS34 *in vitro*^18^, and we and other^48^ found that chemically inhibiting VPS34 inhibits the fission of lysosomal tubules. These two models are not mutually exclusive and could work in synergy, perhaps providing a timed burst of PI(3)P at the site of fission. This could explain why Arf1-PI4KIIIβ positive vesicles are not recruited to all lysosomal tubule fission events. Finally, our results are also consistent with a model where SEC14L2 would transfer phosphatidylinositol (PI) to lysosomes that would be used as a substrate by VPS34 to produce PI(3)P. However, as SEC14L2 binds only weakly to PI compared to PI(3)P^18^, this possibility appears least likely.

The precise role of the PI(3)P signalling contributing to lysosomal tubule fission remains to be elucidated. However, as phosphoinositides are able to recruit specific proteins^52^, it is tempting to speculate that PI(3)P could allow for the recruitment of effectors required for the scission of lysosomal membranes.

Finally, PI(4)P production on lysosomes by PI4KIIIβ was previously proposed to inhibit the formation of lysosomal tubules^13^. We show here that the production of PI(4)P on lysosomes does not impair their ability to tubulate. Instead, we found that the depletion of PI(4)P strongly suppresses the tubulation of lysosomes suggesting that PI(4)P has a pro-tubulation role. This is in line with several studies showing that PI(4)P promotes the formation of tubules in phagolysosomes^14^ and endosomes^15,16,53^. Our results indicate that the extensive tubulation of lysosomes caused by PI4KIIIβ inhibition is the result of impaired fission of lysosomal tubules. We propose that loss of PI4KIIIβ function affects the function and/or the formation of Arf1-PI4KIIIβ positive vesicles since PI4KIIIβ collaborates with Arf1 at the Golgi to promote the formation of vesicles from the TGN^28,54^, thus leading to an increase in lysosome tubule numbers. Moreover, this increased number of lysosomal tubules suggests that PI4KIIIβ does not play a major role in the production of PI(4)P required for lysosomal tubules formation and that type II PI4K, that were proposed to produce PI(4)P on lysosomes^53,55^, are likely generating PI(4)P on lysosomes allowing membrane tubulation to occur. Further studies will be required to determine the source of PI(4)P required for lysosomal tubulation.

In conclusion, we propose that Arf1-PI4KIIIβ positive vesicles contribute to the fission of lysosomal tubules of a wide range of lysosomal organelles by promoting a SEC14L2 dependent PI(3)P signalling at the site of fission.

## Methods

### Antibodies and chemicals

The following antibodies were used in this study: Rabbit anti-LC3B (Novus biologicals, NB100-2220, dilution: 1:1000), Rabbit anti-β-Actin HRP conjugated (Cell signaling, 5125, dilution 1:10 000), rabbit anti-*E. coli* antibodies (Bio-Rad, Bio-Rad, 4329-4096, 1:100), and goat anti-Rabbit HRP (ThermoFisher, 31460, dilution: 1:10000). Brefeldin A (B7651) Invertase (I4504), Rapamycin (R0395), and Sucrose (S0389) were purchased from Sigma. Dynasore and PIK93 were purchased from Tocris. YM201636, ikarugamycin, Bafilomycin A1 and VPS34-IN1 were purchased from Cayman Chemicals. PI4KIIIbeta-IN-10 (HY-100198) was purchased from MedChemExpress. Sheep Red blood cells (10% suspension) and rabbit anti-Sheep antibodies (IgG fraction) were purchased from MP biomedicals.

### Plasmids

The following plasmids were used in this study: GFP-KDEL, CFP-LC3, mCherry-LC3 and ub-RFP-SKL were previously described^56,57^. Lamp1-mCherry was a gift from Amy Palmer (Addgene #45147); Arf1-GFP was a gift from Paul Melancon (Addgene #39514); TGN46-mEmerald was a gift from Michael Davidson (Addgene #54279) and mCherry-FKBP-MTM1 was a gift from Tamas Balla (Addgene #51614); the bacterial expression vector pBAD::mRFP1 was a gift from Robert Campbell (Addgene #54667). GFP-PI4KIIIB, GFP-2xP4M, mCherry-P4M, PX-GFP, GFP-2xFYVE, iRFP-FRB-Rab7, GFP-Rab5, GFP-ORPSAC1 and GFP-EEA1 were a gift from Sergio Grinstein (The Hospital for Sick Children, Toronto). CFP-Rab11 was a gift from John Brumell (The Hospital for Sick Children, Toronto). SEC14L2 human ORF (BC058915) was a gift from Brian Raught (Princess Margaret Cancer Centre, Toronto). Arf1-BFP and Arf1-mCherry were generated by replacing GFP with BFP (from mitoBFP, Gift from William Trimble (The Hospital for Sick Children, Toronto)) and by mCherry (from Lamp1-mCherry), respectively. Arf1-MTM1-mTq2 was generated by inserting MTM1 from mCherry-FKBP-MTM1 in between Arf1 and mTq2 of Arf1-mTq2. SEC14L2-mCherry was made by replacing Lamp1 (in Lamp1-mCherry) with SEC14L2. SEC14L2-GFP was made by replacing Arf1 (in Arf1-GFP) with SEC14L2. SEC14L2-MTM1-GFP was generated by inserting MTM1 from mCherry-FKBP-MTM1 in between SEC14L2 and GFP of SEC14L2-GFP. LysoGFP-Sac1 and LysoGFP-Sac1 C389S were made by subcloning LysoGFP-Sac1 from pLJM1-Lyso-FLAG-GFP-Sac1-Cat-WT (Gift from Roberto Zoncu (Addgene #134645)) and pLJM1-Lyso-FLAG-GFP-Sac1-Cat-CS (Gift from Roberto Zoncu (Addgene #134653)) in a Clontech vector. GFP-ORPSAC1 C392S was generated from GFP-ORPSAC1 by site-directed mutagenesis. Arf1-SNAP, Lamp1-SNAP and PX-SNAP were made by replacing GFP (in Arf1-GFP), mCherry (in Lamp1-mCherry), and GFP (in PX-GFP) with SNAP from SNAP-Sec61b (Gift from Gia Voeltz; Addgene #141152). CFP-GAI-MTM1 was made by inserting MTM1 from mCherry-FKBP-MTM1 after GAI in CFP-GAI(1-92) (gift from Takanari Inoue, Addgene # 37307). LysoYFP-GID1 was made by inserting Lyso from LysoGFP-Sac1 before YFP in YFP-GID1 (gift from Takanari Inoue, Addgene # 37305). zSec14l3-mCherry was a gift from Dr. Shunji Jia (Tsinghua University, Beijing). zSec14l3-ΔCRAL-TRIO-mCherry and zSec14l3-ΔGOLD-mCherry were generated by PCR-mediated deletion of the coding regions of the CRAL-TRIO (amino acid 76 to 249), and the GOLD (amino acid 275 to 384) domains, respectively. zSec14l3-M5 was custom synthesized (GeneScript). DNA sequences were verified by Sanger sequencing.

### siRNAs

The following siRNAs were used: SEC14L2 (siSEC14L2-1: 5’ GCCGAAUCCAGAUGACUAUUU 3’; siSEC14L2-2: 5’ GUGGCCUAUAACCUCAUCAAA 3’; siSEC14L2-3: 5’ CCGAAACACUGAAGCGUCUUU 3’). siRNAs were custom synthesized from Millipore Sigma, and MISSION® siRNA Universal Negative Control #1 (Sigma) was used as a negative control.

### Cell culture and transfection

COS-7, HeLa, MEFs, HEK293 and RAW 264.7 cells were obtained from ATCC and were cultured in DMEM (Gibco) supplemented with 10% FBS (Wisent) at 37°C in humidified air containing 5% CO2. Primary mouse macrophages were obtained and cultured as previously described^8^. Transfection was performed using the Neon transfection system (Invitrogen) according to the manufacturer’s instructions, using the following parameters: 1050V, 30ms and two pulses (COS-7); 1005V, 35ms and two pulses (HeLa); 1350V, and 30ms and one pulse (MEFs), 1150v 20ms and two pulses (HEK 293) and 1680v 20ms and one pulse (RAW 264.7). 5µg of DNA for 0.5 × 10^6^ cells (1 × 10^6^ cells for RAW 264.7) was used, and cells were imaged or treated 24 hours after transfection. A siRNA final concentration of 100nM was used for silencing experiments, and cells were imaged or treated 48 hours after transfection.

### Fixed-cell imaging of phagosome resolution

RAW 264.7 cells were seeded at 20% confluence on #1.5 coverglass (Electron Microscopy Sciences) the day prior to phagosome fragmentation. Following the drug treatment and phagosome resolution, cells were fixed with 4% paraformaldehyde (Electron Microscopy Sciences) for 15 minutes, then washed 3 times with PBS. External bacteria were labeled with rabbit anti-E. coli antibody followed by DyLight 488-conjugated anti-rabbit antibodies. Cells were then mounted on a slide using Dako Mounting Medium (Agilent). Images were acquired on a Quorum spinning disk system (Quorum, Guelph, ON) consisting of a Leica DMi8 microscope (Leica), an Andor Diskovery Multimodal Imaging system (Oxford Instruments), and an Andor Zyla 4.2 Megapixel sCMOS camera (Oxford Instruments), and controlled by Quorum Wave FX Powered by MetaMorph software (Quorum).

### Live imaging

Most images were acquired on a Leica SP8 laser scanning Confocal microscope equipped with a Lightning (deconvolution) module. Cells were grown in Lab-Tek^TM^ chambers (Nunc) with a borosilicate glass bottom. Prior imaging, media was replaced, and cells were imaged using a Leica SP8 with a 63x glycerol immersion objective lens, 1.3 NA (Leica) in a temperature (37°C) and atmosphere (5% CO2) controlled environment. Depending on the experiments, a single optical section or a Z-stack of 3 to 5 steps with a step size ranging from 250nm to 340nm was acquired. The Las X software (Leica) was used to perform acquisition, and images were deconvoluted with the lightning module (Leica) using adaptative parameters. For lysosomal tubule fission events, a video of 1.5 minutes with 1 frame every 2 seconds was acquired.

Some images were acquired on a Zeiss LSM980 laser scanning Confocal microscope equipped with an Airyscan 2 module, with a 63x oil immersion objective lens, 1.3 NA (Zeiss) in a temperature (37°C) and atmosphere (5% CO2) controlled environment, in which case this is indicated in the text and/or legends. Cells imaged were of similar low intensity in order to be able to compare cells with similar levels of overexpression.

### Lattice Light Sheet microscopy (LLSM)

A ZEISS Lattice Lightsheet 7 system was used for this study. Samples were grown and imaged in Lab-Tek^TM^ chambers (Nunc) in a temperature (37°C), CO_2_ (5%) and humidity (70%) controlled environment. Samples were illuminated by optical light sheets generated by beam shaping by cylinder lens and spatial light modulator (SLM) with beams of 30µm in length and 1000nm of thickness using 488nm and 561nm lasers and appropriate filters. Fluorescent emission was collected by a detection objective (lens 44.83× / 1.0 (at 60° angle to cover glass)) and a Pco.edge 4.2 CLHS sCMOS camera (6.5 um pixel size, 2048×2048, 82% QE). Acquired data were deconvoluted (Constrained iteration using automatic strength) and deskewed using Zen 3.3 software (Zeiss).

### Three-dimensional reconstruction

three-dimensional reconstructions of fluorescent images were performed using Imaris Bitplane (9.2.1) using the “Make surface” feature with default parameters.

### Gibberellin-induced dimerization system

For Gibberellin-induced dimerization experiments, GID1 tagged membrane anchor proteins were used to recruit GAI fused cytosolic construct upon GA_3_-AM treatment (10µM). Cells were imaged after 1 hour of GA_3_-AM treatment unless indicated in the text and/or legend.

### Image Analysis

The number of lysosomal tubules per cell was counted manually using ImageJ software, normalized to the volume or area of cells and expressed as “Number of Lamp1 tubules/µm^3^” or “Number of Lamp1 tubules/µm^2”^. For lysosomes-Arf1 positive vesicle contact evaluation, the number of lysosomes showing contact with at least one Arf1 vesicle at one time frame was counted and was then expressed as a percentage. The minimum duration of contacts between these two compartments was analyzed as previously described^19^. Briefly, 10 randomly chosen contact were analyzed. Only contacts that were already formed at the beginning of the acquisition and that lasted for at least 3 consecutive frames (>6 s) were analyzed. The minimum duration of contact was quantified as the time before the dissociation of the lysosome-Golgi-derived vesicle contact over a 1.5 min video. Contacts that lasted during the totality of the video were scored as 1.5 min (90s) contacts.

Lamp1 positive organelles and Lysosomal tubule fission events were defined as events showing a clear division of a tubule from one Lamp1 positive organelle or from one parent lysosome organelle (Autolysosome, Endolysosome or Phagolysosome). In the vast majority of the cases (>90%), the fission event was observed at the tubule neck. However, fission events occurring in the middle of the tubules were also counted as positive. The presence of vesicle markers such as Arf1-GFP at the tubule fission site was defined as the contact of the vesicle with the tubule fission site at the frame just before the fission event was observed. The presence of vesicle markers at the fission site was then expressed as a percentage. To compare the percentage of fission sites marked by these vesicle markers to that of one expected by chance, we performed the same analysis, but after the Lamp1-mCherry channel was rotated by 90°. For the rate of Lamp1 tubule fission, the number of Lamp1 tubule fission events was counted and then normalized by time and volume to calculate a fission rate (Normalized rate of Lamp1 tubule fission).

For phagosome resolution/fragmentation, images were converted to binary images. The intensity threshold for binary conversion was selected manually to preserve as many vesicular structures as possible while minimizing the merging of objects after binary conversion. This was then applied to all images across a biological replicate. When necessary, the watershed function was used to further segment particles on the binary images. Phagosome fragments were counted using ImageJ’s “Analyze particles” function. Fragments were defined as objects with an area between 10 and 1000 pixels (0.04 to 3.83 µm^2^. Intact phagosomes and external particles were manually subtracted from the count. Fragments from whole cells were included in the analysis, while we excluded regions where cells were overly crowded, making dissection difficult. Fragments per cell were then quantified.

To estimate the levels of PI(3)P at lysosomes, the PI(3)P probe PX^46^ was used. A mask of lysosomes (using cresyl violet or overnight chased 10kDa Dextran 647) was generated using an automatic threshold of ImageJ, and the mean fluorescent intensity of PX was measured. This was normalized to the cytosolic levels of PX. To evaluate whether the fission of Lamp1 tubules was associated with a regulation of PI(3)P and PI(4)P, we identified Lamp1 tubules fission events and measured these phosphoinositides using appropriate fluorescent biosensors (PX and 2xFYVE for PI(3)P; 2xP4M for PI(4)P) for 8 frames (16 seconds) before the fission event and 3 frames (6 seconds) after it was observed. The fluorescence intensity was measured on the Lamp1 positive organelle main body and tubule neck only. The mean fluorescence intensity of the biosensor was normalized to that of Lamp1-mCherry for each respective frame.

Enlarged autolysosomes were defined as Lamp1 and LC3 positive structures of more than 1µm in diameter.

The number of peroxisomes per cell was counted using Volocity software (Perkin Elmer) and was normalized to the volume of the cells.

Whenever possible, we performed a blind analysis of our data. After acquisition, files were anonymized before analysis.

### Quantitative PCR and primers

The Monarch® Total RNA Miniprep Kit (New England BioLabs) was used to perform total cellular RNAs isolation, following manufacturer instructions, and cDNAs were subsequently synthesized using the High-Capacity cDNA Reverse Transcription kit (Applied Biosystems). qPCR was performed using a StepOne Real Time PCR System using SYBR green (ThermoFisher Scientific) using β-actin as a reference gene for all quantifications.

### Induced lysosomal tubule fission

Lysosomal tubule fission was promoted using four different ways: (1) by incubating cells for 8h in an amino acid-free media HBSS (Hanks’ Balanced Salt Solution) that triggers autophagic lysosome reformation which implicates formation of tubules from autolysosomes and their fission; (2) by reversibly inducing lysosomal swelling by incubating cells with 30mM sucrose for 24 hours, that leads to the formation of sucrosomes (Swollen lysosomes full of sucrose). Mammalian cells cannot degrade sucrose which builds up in lysosomes leading to the formation of sucrosomes. Cells were then incubated with Invertase (0.5mg/mL) for 1 hour, allowing its endocytosis and delivery to lysosomes leading to sucrose degradation and subsequent lysosomal shrinkage in a process involving strong lysosomal fission; (3) by inducing phagolysosomes formation in RAW 264.7 macrophages by incubating cells with opsonized SRBCs and (4) by reversibly inhibiting PIKFyve with YM201636 (1µM for 1-hour treatment) which leads to extensive lysosomal swelling. Upon washout of the drug (two subsequent washing with regular culture media), lysosomes shrink to come back to their normal size in a process that involves strong lysosomal fission.

### Phagocytosis

For phagocytosis experiments (except for phagosome fragmentation assay), opsonized sheep red blood cells (SRBCs) were used as prey for RAW264.7 macrophages. For opsonization; 100µL of 20% SRBCs suspension was washed with PBS and opsonized with 3µL of rabbit IgG fraction against SRBC for 1 hour at 37°C under agitation. SRBCs were then washed twice with PBS and resuspended in 1000µL of PBS. Phagocytosis was induced by incubating cells with a 100-fold dilution of opsonized SRBCs suspension in regular media. Phagocytosis was synchronized by centrifuging the imaging chamber for 10s at 300g. Cells were then washed with HBSS and were imaged in HBSS.

### Phagosome fragmentation assay

The assay was performed as previously described^8^. In brief, fluorescently labeled E. coli (DH5α E. coli strain) were transformed with the pBAD::mRFP1 plasmid^58^, and RAW 264.7 were allowed to internalize mRFP1-labeled E. coli. Cells were then washed with regular media and incubated for the indicated time. Cells were treated with either DMSO, 10 µM PIK93, 10 µM Brefeldin A (BFA), or 0.5 µg/mL Ikarugamycin (Ika) 1 hour after phagocytosis and imaged after 5 hours of treatment using a spinning disk confocal microscope.

### Western blotting

Cells were lysed in a Lysis buffer (100 mM Tris-HCL, 1% SDS) containing a protease inhibitor cocktail (BioShop). Lysates were then incubated for 15 min at 95°C and vortexed briefly every 5 min during this incubation. Lysates were then centrifugated at 21,000×g for 15 min at room temperature. Supernatant was then collected, and their protein concentration was determined using the BCA assay kit (Thermo Scientific). A ChemiDoc (Bio-Rad Laboratories) system was used to visualize proteins.

### Labelling of lysosomes with Cresyl violet and fluorescent Dextran

Cresyl Violet (Sigma-Aldrich) staining was used to label acidic lysosomes^23^. Cells were incubated with 1µM of Cresyl violet in HBSS for 3 minutes at 37°C, were then rinsed twice and incubated in appropriate media for imaging.

Alternatively, lysosomes were also stained using fluorescent 10 kDa Dextran A647 (D22914; Thermo Fisher). Cells were incubated with 0.1mg/mL of Dextran for 3 hours at 37°C. Cells were then rinsed twice and chased overnight in appropriate media to allow the Dextran to reach lysosomes before imaging.

### SNAP-tag labelling

To label SNAP tag-containing constructs, cells were incubated for 30 min at 37°C in appropriate media containing 3µM of cell-permeable SNAP-Cell® 647-SiR (New England BioLabs). Cells were then rinsed twice with fresh media and incubated for 30min at 37°C. Media was then replaced, and cells were imaged.

### Graphing and figure assembly

Graphics were prepared using GraphPad Prism 5 or 9.3. The brightness and contrast of microscopy images were adjusted using ImageJ software for presentation purposes. For the experiment with constructs exhibiting a strong cytosolic signal (Arf1-GFP, Arf1-BFP, Arf1-SNAP, PX-SNAP and GFP-PI4KIIIB), brightness and contrast were adjusted to mask the cytosolic signal allowing better visualization of vesicles, for presentation purposes. All final figures were assembled using Illustrator CS6 (Adobe).

### Statistics and Reproducibility

All statistical tests were performed using GraphPad Prism 5 and 9 and are described in the figure legends. A p-value of p<0.05 was considered statistically significant. All Graphs show the mean ± SEM from three independent trials (unless specified in the legends).

## Data availability

All data that support the findings of this study are included in the manuscript or are available from the authors upon reasonable request.

## Acknowledgments

We thank Sharon Leung for critically reading the manuscript; Drs. Sergio Grinstein, Spencer Freeman and John Brumell for providing reagents; Drs. Paul Paroutis and Kimberly Lau at the SickKids Imaging Facility for assistance with live-cell imaging. Infrastructure for the Kim Laboratory was provided by a John Evans Leadership Fund grant from the Canadian Foundation for Innovation and the Ontario Innovation Trust. This work was supported by operating grants from the Canadian Institutes of Health Research to P.K.K (PJT#180476) and R.J.B (PJT-166047 and PJT-183914) And Canada Research Chairs to R.J.B. (950-232333); M.B was supported by a Restracomp Fellowship from the Hospital for Sick Children and a CIHR Postdoctoral fellowship from the Canadian Institutes of Health Research.

## Author contributions

M.B. designed the experiments, performed most of the experiments and analyzed the data. L.F.D, N.D, A.F and S.M performed imaging experiments and analyzed the data. R.J.B. designed and analyzed the macrophage experiments. P.K.K. supervised and coordinated the overall research and analyzed the data. M.B. and P.K.K. wrote the manuscript.

## Supplementary Figures legends

**Supplementary figure 1.**
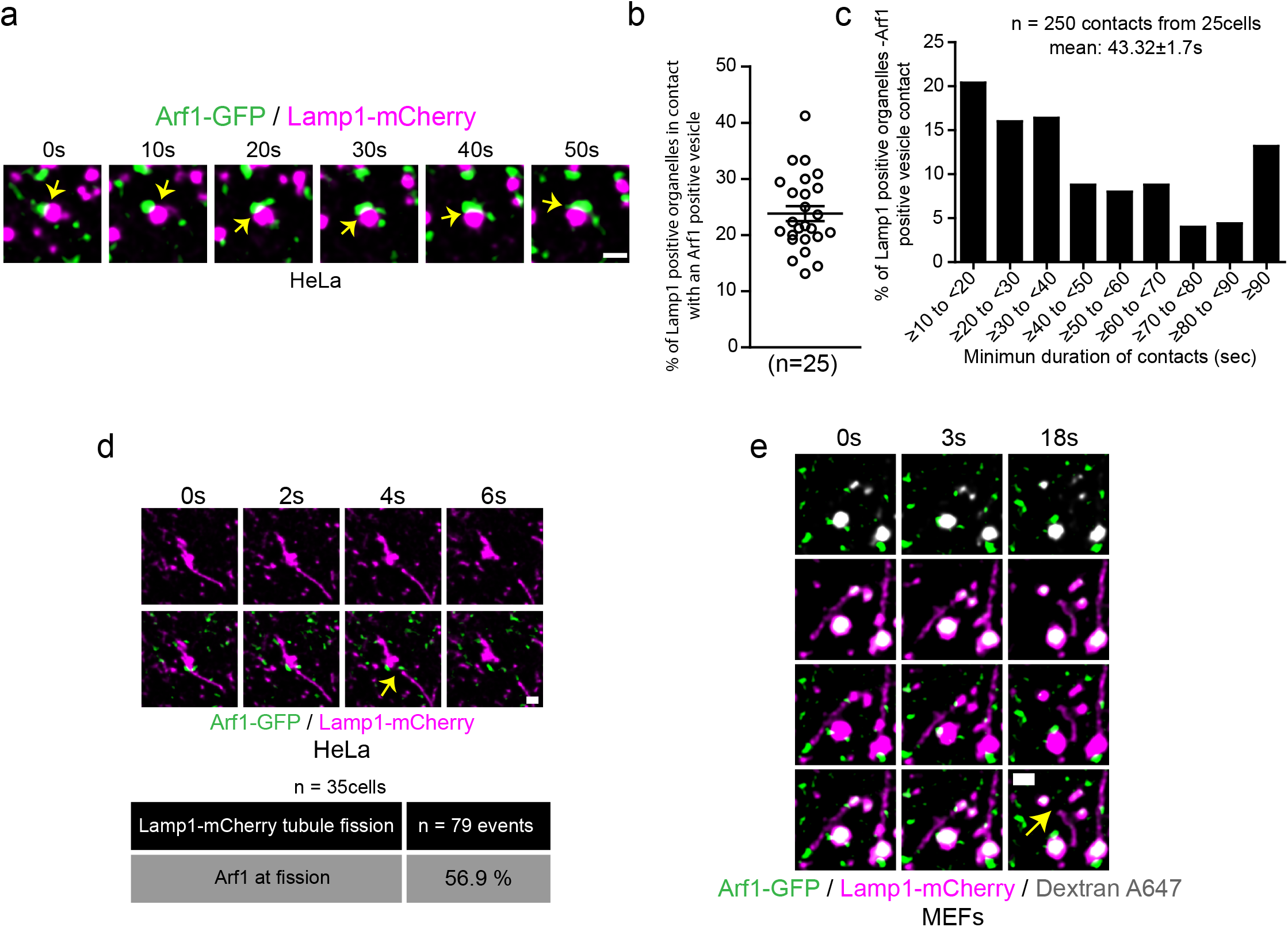
Arf1 positive vesicles make stable and dynamic contacts with Lamp1 positive organelles in HeLa cells and mark lysosomal tubule fission sites. **a**. Representative time-lapse live images showing dynamic contact between Arf1-GFP positive vesicles and Lamp1 positive organelles in HeLa cells. Yellow arrow points to contacts between these two compartments. Scale bar: 1µm. **b**. Quantification of the percentage of Lamp1 positive organelles in contact with at least one Arf1 positive vesicle in HeLa cells. The graphs show the mean ± SEM, cells from three independent experiments. **c**. Quantification of the mean minimum duration (seconds) of Lamp1 positive organelle-Arf1 positive vesicle contacts. A contact is defined by overlapping pixels, and the time of contact was quantified from time-lapse images taken at 1 frame every 2 seconds. 250 randomly chosen contacts from 25 HeLa cells were analyzed. **d**. Representative time-lapse imaging showing an Arf1-GFP positive vesicle marking the fission site of tubule from a Lamp1-mCherry positive organelle in a HeLa cell starved for 8h with HBSS (amino acid-free media) in order to promote formation and fission of Lamp1 tubules. Yellow arrow indicates the fission event. Scale bar = 1µm. The percentage of tubule fission events marked by Arf1-GFP vesicles was quantified. n=79 events from 35 HeLa cells. **e**. Representative time-lapse imaging showing an Arf1-GFP vesicle marking fission site of a tubule from a lysosome in a MEF cell. Lysosomes were identified as organelles positive for Lamp1 and overnight chased fluorescent 10kDa Dextran. Yellow arrow indicates fission. Scale bar = 1µm.

**Supplementary figure 2.**
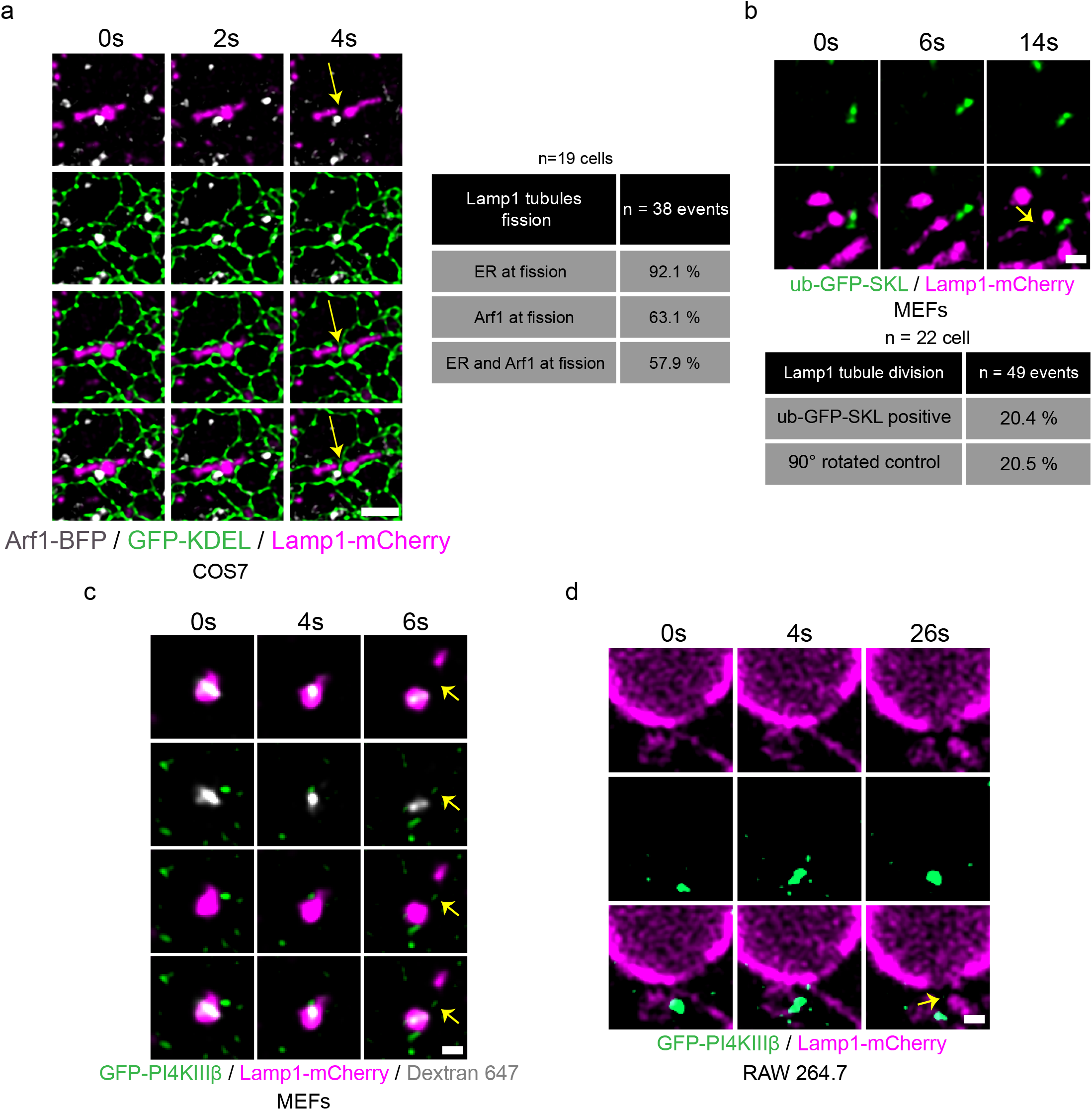
The ER is present at Lamp1 tubule fission sites marked by Arf1 positive vesicles and PI4KIIIβ positive vesicles are found at endolysosomal and phagolysosomal tubule fission sites. **a**. Time-lapse imaging of a COS7 cell expressing the ER marker GFP-KDEL, Arf1-BFP and Lamp1-mCherry incubated for 8h in HBSS media that shows the concomitant presence of the ER and of an Arf1 positive vesicle to a Lamp1 positive tubule fission event. Quantification of such event are shown in the table. n=38 events from 19 cells. Yellow arrow indicates fission. Scale bar: 2µm. **b**. Representative time-lapse images showing the absence of ub-GFP-SKL (Peroxisome) at a Lamp1 positive tubule fission site in MEFs starved (HBSS) for 8 hours and quantification of percentage of fission events marked by ub-GFP-SKL. Negative control analysis was performed with the Lamp1-mCherry signal rotated by 90°. n = 49 events from 22 cells. Yellow arrow indicates fission. Scale bar = 1µm. **c-d**. Representative time-lapse imaging showing a GFP-PI4KIIIβ positive vesicle marking fission site of a tubule from an (**c**) endolysosome in a MEF cell. Endolysosomes were formed by 24 hours incubation with sucrose and their tubulation and fission was observed after 1 hour of treatment with invertase (0.5mg/mL). Endolysosomes were defined as positive for Lamp1 and overnight chased fluorescent 10kDa Dextran; (**d**) Phagolysosome in RAW264.7 cells phagocyting SRBCs. Yellow arrows indicate fission. Scale bar = 1µm.

**Supplementary figure 3.**
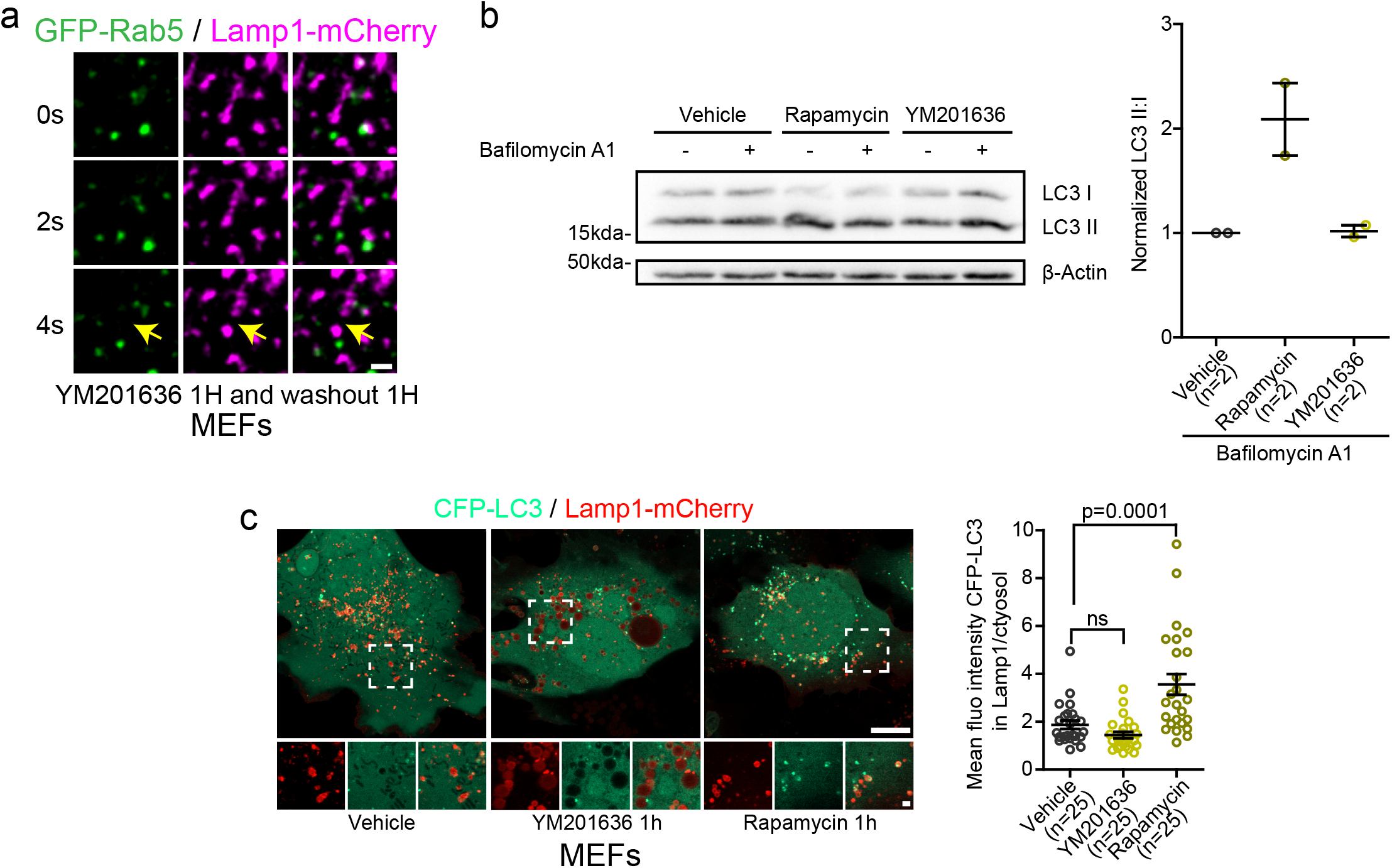
Short-term inhibition of PIKfyve with YM201636 does not induce autophagy or formation of endolysosomes. **a**. Time-lapse imaging of a Lamp1 positive tubule fission event in a MEF cell expressing Rab5-GFP and Lamp1-mCherry after short-term inhibition of PIKfyve using YM2013636 (1µM for 1 hour) followed by 1 hour washout of the drug. The Yellow arrow indicates tubule fission. Scale bar = 1 µm. **b**. Representative western blot images of MEFs treated 1 hour with YM201636 (1µM) or vehicle control, rapamycin (10µg/mL) was used as a positive control. Bafilomycin A1 treatment (500nM, 1 hour) was used to inhibit degradation of autophagosomes allowing to evaluate whether YM201636 treatment induced formation of autophagosomes. Levels of LC3II were normalized to LC3I to quantify autophagosomes. β-Actin was used as a positive control. The graphs show the mean ± SEM, cells from two independent experiments. **c**. Representative images of MEFs cells expression CFP-LC3 and Lamp1-mCherry and treated with YM201636 (1µM for 1h) or Rapamycin (10µg/mL) as a positive control. Scale bars: 10µm and 2µm (inset). Quantification of the mean fluorescence intensity of CFP-LC3 colocalizing with Lamp1 normalized to that of the cytosol. The graphs show the mean ± SEM, cells from three independent experiments. One-way ANOVA with Dunnett’s Multiple Comparison Test. ns = 0.4570.

**Supplementary figure 4.**
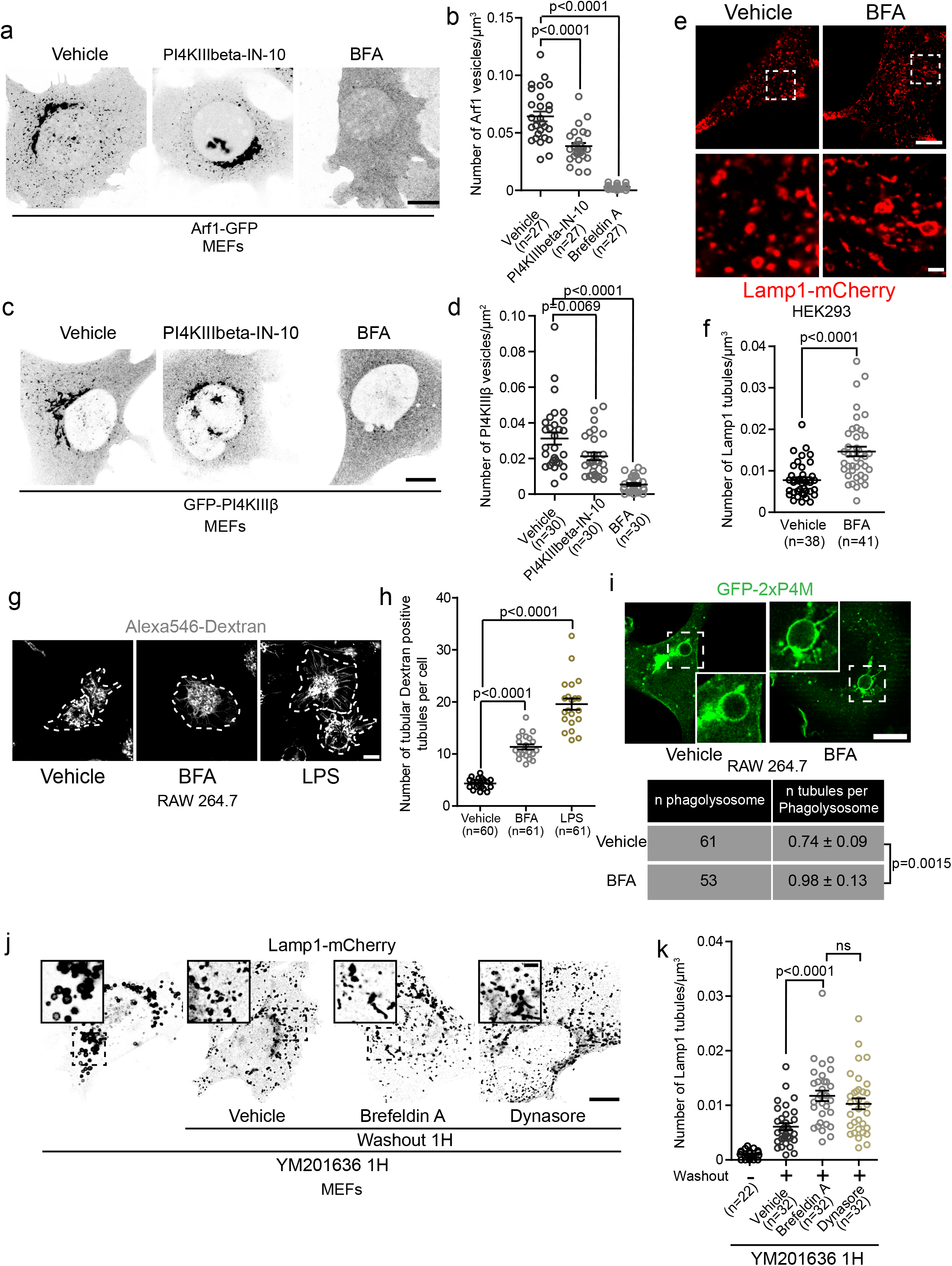
Inhibition of formation and/or function of Arf1-PI4KIIIβ positive vesicles increases the number of lysosomal tubules. **a-b**. (**a**) Representative images of MEFs expressing Arf1-GFP and treated with PI4KIIIbeta-IN-10 (25nM for 3h) or BFA (10µg/mL for 1h). Ethanol was used as a vehicle control. Scale bar: 10µm. (**b**) Quantification of the number of Arf1-GFP positive vesicles per cell. One-way ANOVA with Dunnett’s Multiple Comparison Test. **c-d**. (**c**) Representative images of MEFs expressing GFP-PI4KIIIβ and treated with PI4KIIIbeta-IN-10 (25nM for 3h) or BFA (10µg/mL for 1h). Ethanol was used as a vehicle control. Scale bar: 10µm. (**d**) Quantification of the number of GFP-PI4KIIIβ positive vesicles per cell. One-way ANOVA with Dunnett’s Multiple Comparison Test. **e-f**. (**e**) Representative images of HEK293 cells expressing Lamp1-mCherry and treated with BFA (10µg/mL for 1h) or Ethanol as a vehicle control. Scale bars: 10µm and 1µm (inset) (**f**) Quantification of the number of Lamp1 positive tubules in these cells. Two-sided unpaired t-test. **g-h**. (**g**) Representative images of primary mouse macrophages that were pulsed with Alexa546-Dextran (50µg/mL for 30min) to mark lysosomes and treated with BFA (5µg/mL for 2h). LPS (500ng/mL for 2h) was used as a positive control. DMSO was used as a negative control (vehicle). Images were acquired using a Spinning disk confocal microscope system (Quorum Technologies). White dotted lines show cell outline. Scale bar: 10µm. (**h**) Quantification of the number of Dextran positive tubules in these cells. One-way ANOVA with Dunnett’s Multiple Comparison Test. **i**. Representative images of phagolysosomes from RAW264.7 macrophages expressing GFP-2xP4M and treated with BFA (10µg/mL) 15 minutes after induction of phagocytosis of SRBCs. Cells were imaged from 15min to 30min after BFA was added. Ethanol was used as vehicle. Scale bar: 10µm. Quantification of the number of tubules from these cells. Two-sided unpaired t-test. **j-k**. (**j**) Representative images of MEFs expressing Lamp1-mCherry. Cells were treated with YM201636 (1µM for 1h) to inhibit PIKfyve and were then washout to remove PIKfyve inhibitor in presence of BFA (10µg/mL) or Dynasore (40µM) and imaged 1h after. Scale bars: 10µm and 2µm (inset) (**k**) Quantification of the number of Lamp1 positive tubules from these cells. One-way ANOVA with Dunnett’s Multiple Comparison Test. All graphs (**b, d, f, h, k**) show the mean ± SEM, cells from three independent experiments.

**Supplementary figure 5.**
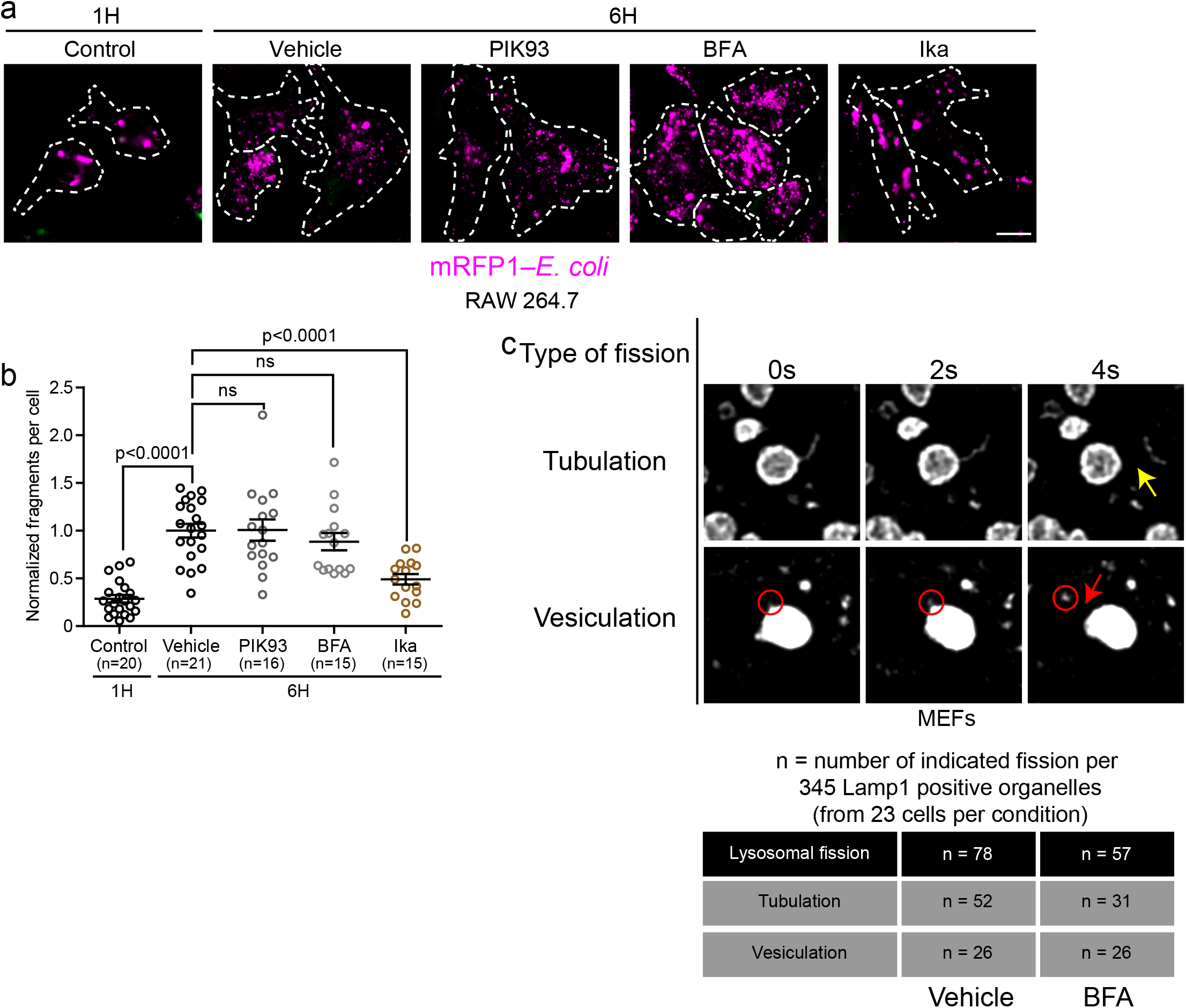
Inhibition of PI4KIIIβ or Arf1 activation does not affect lysosomal fission by vesiculation and splitting. **a-b**. (**a**) Representative images of phagosome fragmentation in RAW 264.7 that were allowed to internalize mRFP1-labeled *E. coli* (magenta), then washed and incubated for 1 hour or 6 hours before imaging. Cells were treated with either DMSO, 10 µM PIK93, 10 µM Brefeldin A (BFA), or 0.5 µg/mL Ikarugamycin (Ika) 1 hour after phagocytosis and imaged at indicated times. Cells were labeled with anti-*E. coli* (green) to identify external bacteria. Images were acquired using a Spinning disk confocal microscope system (Quorum Technologies). Scale bar = 10 µm. (**b**) Quantification of the number of mRFP-*E.coli* positive fragments in these cells. The graphs show the mean ± SEM, cells from three independent experiments. Statistical analysis was performed using the Kruskal-Wallis test with Dunn’s multiple comparisons test. ns: p>0.9999 (Vehicle vs PIK93) and p=0.8132 (Vehicle vs BFA). **c**. MEFs cells expressing Lamp1-mCherry were treated with the PIKfyve inhibitor YM201636 (1µM for 1h) and then PIKfyve inhibitor was washed out in presence of BFA (10µg/mL) or a vehicle control (ethanol). Cells were imaged between 5 and 30 min after washout. A total of 345 Lamp1 positive organelles (from 23 cells) per condition were analyzed for two types of lysosomal fission: fission of tubules (tubulation) or of vesicles (vesiculation). The Yellow arrow indicates tubule fission while the red circle indicates a vesiculation event and the red arrow the fission of this vesicle from the lysosome. Scale bar: 1µm.

**Supplementary figure 6.**
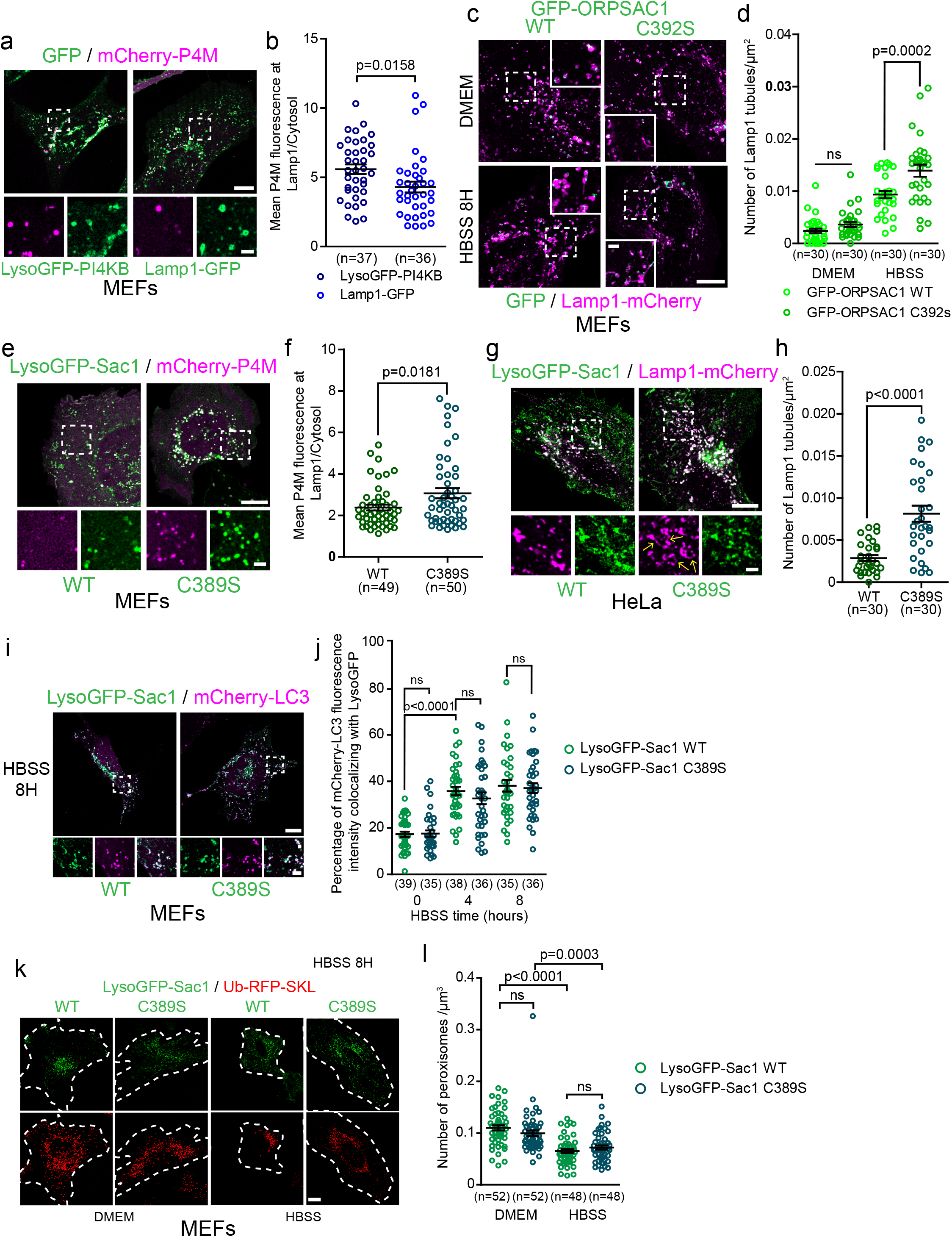
Depletion of lysosomal PI(4)P impairs formation of Lamp1 positive tubules but does not affect the fusion of lysosomes with autophagosomes. **a-b.** (**a**) Representative images of MEFs expressing mCherry-P4M and LysoGFP-PI4KB or Lamp1-GFP. Scale bars: 10µm and 2µm (inset). (**b**) Quantification of the P4M fluorescence intensity in Lamp1 mask normalized by the cytosolic one. Two-sided unpaired t-test. **c-d.** (**c**) Representative images of MEFs cells expressing Lamp1-mCherry or GFP-ORPSAC1 to deplete lysosomal PI(4)P. The catalytic dead version of ORPSAC1 (C392S) was used as a negative control. Cells were treated with HBSS for 8h to promote formation of tubules from Lamp1 positive organelles. Scale bars: 10µm and 2µm (inset). (**d**) Quantification of the number of Lamp1 positive tubules for these cells. Two-way ANOVA with Tukey’s multiple comparisons test. ns: 0.6594. **e-f.** (**e**) Representative images of MEFs expressing mCherry-P4M and LysoGFP-Sac1 or the catalytic dead version of it (C389S). Scale bars: 10µm and 2µm (inset). (**f**) Quantification of the P4M fluorescence intensity in Lamp1 mask normalized by the cytosolic one. Two-sided unpaired t-test. **g-h.** (**g**) Representative HeLa cells expressing Lamp1-mCherry and LysoGFP-Sac1 or the catalytic dead version of it (C389S). Cells were treated with HBSS to promote formation of tubules from autolysosomes. Yellow arrow in the inset point to Lamp1 tubules. Scale bars: 10µm and 2µm (inset). (**h**) Quantification of the number of tubules from these cells. Two-sided unpaired t-test. **i-j.** (**i**) Representative images of MEFs cells expressing mCherry-LC3 and LysoGFP-Sac1 or the catalytic dead version of it (C389S). Cells were treated with HBSS to promote formation of autophagosomes and their fusion with lysosomes. Scale bars: 10µm and 2µm (inset). (**j**) Quantification of the percentage of mCherry-LC3 staining colocalizing with LysoGFP at indicated time of HBSS treatment. Two-way ANOVA with Tukey’s multiple comparisons test. ns: p>0.9999 (0h), p=0.9456 (4h) and p>0.9999 (8h). **k-l.** (**k**) Representative images of MEFs cells expressing the peroxisomal marker ub-RFP-SKL and LysoGFP-Sac1 or the catalytic dead version of it (C389S). Cells were treated with HBSS for 24h to promote removal of peroxisomes by the autophagic pathway. Scale bar: 10µm. (l) Quantification of the number of peroxisomes per area for these cells. Two-way ANOVA with Tukey’s multiple comparisons test. ns: p=0.3683 and p=0.7442. **b, d, f, h, j, l.**. All graphs show the mean ± SEM, cells from three independent experiments.

**Supplementary figure 7.**
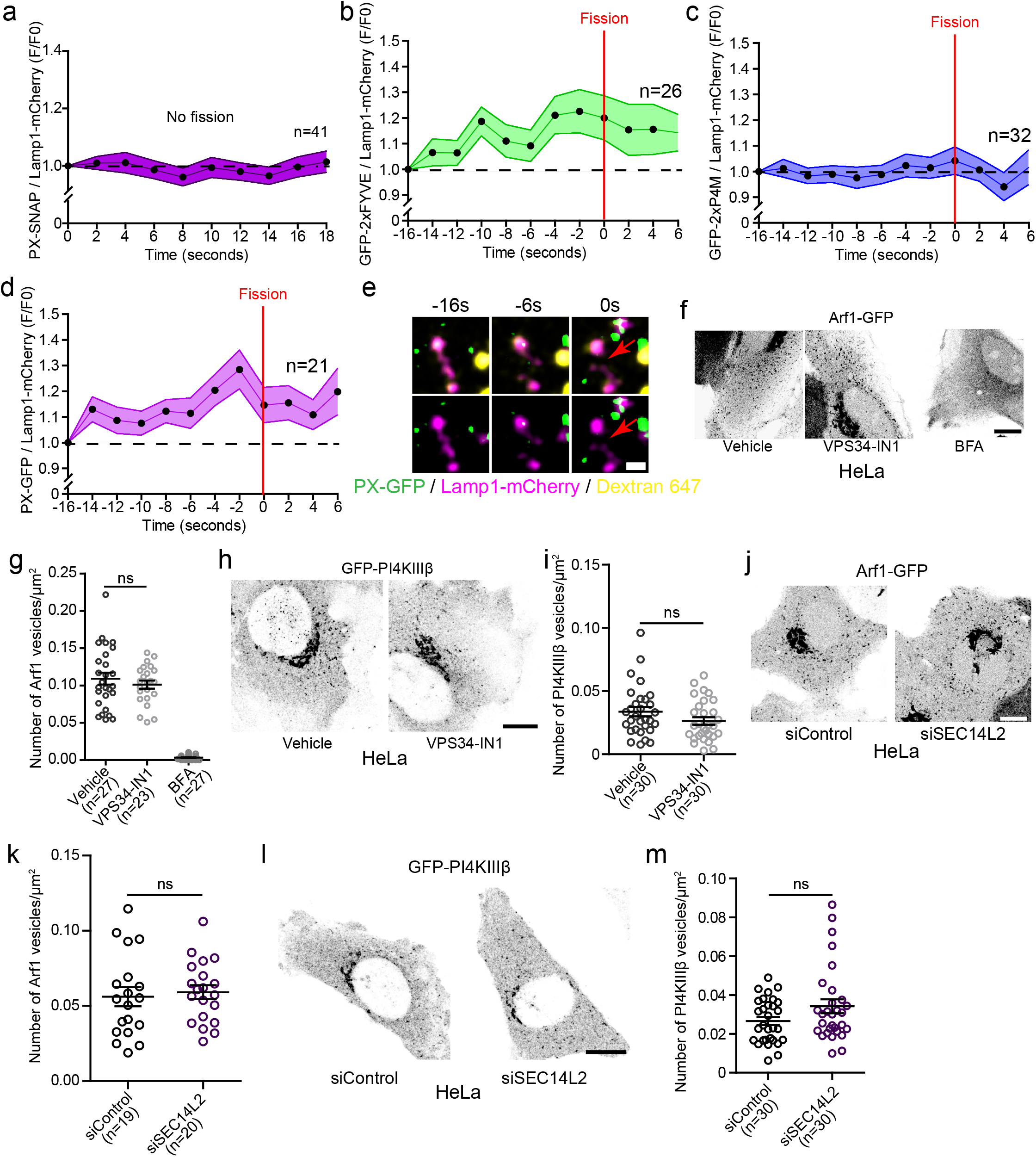
PI(3)P and PI(4)P levels during Lamp1 positive tubule fission events and SEC14L2 depletion or VPS34-IN1 do not affect Arf1-PI4KIIIβ positive vesicles formation. **a**. Normalized fluorescence intensity of PX (PI(3)P biosensor) at Lamp1 positive organelles with tubules that are nor undergoing any fission events. MEFs were starved (HBSS) for 8H to promote formation of tubules from Lamp1 positive organelles. **b**. Normalized fluorescence intensity of 2xFYVE (PI(3)P biosensor) at Lamp1 positive organelles during tubule fission events. MEFs were starved (HBSS) for 8H. **c**. Normalized fluorescence intensity of 2xP4M (PI(4)P biosensor) at Lamp1 positive organelles during tubule fission events. MEFs cells were starved (HBSS) for 8H. **d**. Normalized fluorescence intensity of PX (PI(3)P biosensor)at lysosomes (Lamp1 and overnight chased fluorescent 10 kDa Dextran). MEFs were starved (HBSS) for 8H. Red arrows indicate fission. Scale bar: 1µm. **f-g**. (**f**) Representative images of HeLa cells expressing Arf1-GFP and treated with VPS34-IN1 (1µM) or BFA (10µg/10mL) for 1 hour. Scale bar: 10µm. (**g**) Quantification of the number of Arf1-GFP positive vesicles per cell. One-way ANOVA with Dunnett’s multiple comparison test. ns = 0.5088. **h-i**. (**h**) Representative images of HeLa cells expressing GFP-PI4KIIIβ and treated with the indicated drug for 1H. Ethanol was used as a Vehicle control. Scale bar: 10µm. (**f**) Quantification of the number of GFP-PI4KIIIβ positive vesicles in these cells. Two-sided unpaired t-test. ns = 0.1194. **j-k**. (**j**) Representative Airyscan images of HeLa cells expressing Arf1-GFP and treated with indicated siRNAs. Scale bar: 10µm. (**k**) Quantification of the number of Arf1-GFP positive vesicles per cell. Two-sided unpaired t-test. ns = 0.6999. **l-m**. (**l**) Representative images of HeLa cells expressing GFP-PI4KIIIβ and treated with the indicated siRNAs. (**m**) Quantification of the number of GFP-PI4KIIIβ positive vesicles in these cells. Two-sided unpaired t-test. ns = 0.0676. **a-d, g, i, k, m.** All graphs show the mean ± SEM, cells from two independent experiments.

## Description of Additional Supplementary Files

**File Name: Supplementary Movie 1**

Description; Arf1 positive vesicles and Lamp1 positive organelles appear to form dynamic contacts Movie relative to Figure 1c: Representative time lapse images of a Lamp1 positive organelle-Arf1 positive vesicle contact. MEF cell expressing Arf1-GFP and Lamp1-mCherry.

**File Name: Supplementary Movie 2**

Description: Arf1 positive vesicles are recruited to sites of Lamp1 positive tubule fission Movie relative to Figure 1f: Representative time lapse images of a Lamp1 positive tubule fission event showing recruitment of a Arf1 positive vesicle at the site of fission. MEF cell expressing Arf1-GFP and Lamp1-mCherry.

**File Name: Supplementary Movie 3**

Description: Arf1 positive vesicles are recruited to sites of autolysosomal tubule fission Movie relative to Figure 3b: Representative time lapse images of an autolysosomal tubule fission event showing recruitment of a Arf1 positive vesicle at the site of fission. COS7 cell expressing Arf1-GFP, Lamp1-SNAP(647) and mCherry-LC3 and incubated in amino acid free media (HBSS) for 8 hours.

**File Name: Supplementary Movie 4**

Description: Arf1 positive vesicles are recruited to sites of phagolysosomal tubule fission Movie relative to Figure 3f: Representative time lapse images of phagolysosomal tubule fission event showing recruitment of a Arf1 positive vesicle at the site of fission. RAW 264.7 cells expressing Arf1-GFP and Lamp1-mCherry undergoing phagocytosis of opsonized SRBCs (Sheep Red Blood Cells).

